# Microhomology-mediated end joining drives complex rearrangements and over-expression of *MYC* and *PVT1* in multiple myeloma

**DOI:** 10.1101/515106

**Authors:** Aneta Mikulasova, Cody Ashby, Ruslana G. Tytarenko, Pingping Qu, Adam Rosenthal, Judith A. Dent, Katie R. Ryan, Michael A. Bauer, Christopher P. Wardell, Antje Hoering, Konstantinos Mavrommatis, Matthew Trotter, Shayu Deshpande, Shmuel Yaccoby, Erming Tian, Jonathan Keats, Daniel Auclair, Graham H. Jackson, Faith E. Davies, Anjan Thakurta, Gareth J. Morgan, Brian A. Walker

## Abstract

*MYC* is a widely acting transcription factor and its deregulation is a crucial event in many human cancers. *MYC* is important biologically and clinically in multiple myeloma, but the mechanisms underlying its dysregulation are poorly understood. We show that *MYC* rearrangements are present in 36.0% of newly diagnosed myeloma patients, as detected in the largest set of next generation sequencing data to date (n=1267). Rearrangements were complex and associated with increased expression of *MYC* and *PVT1*, but not other genes at 8q24. The highest effect on gene expression was detected in cases where the *MYC* locus is juxtaposed next to super-enhancers associated with genes such as *IGH*, *IGK*, *IGL*, *TXNDC5*/*BMP6*, *FAM46C* and *FOXO3*. We identified three hotspots of recombination at 8q24, one of which is enriched for *IGH-MYC* translocations. Breakpoint analysis indicates primary myeloma rearrangements involving the *IGH* locus occur through non-homologous end joining, whereas secondary *MYC* rearrangements occur through microhomology-mediated end joining. This mechanism is different to lymphomas, where non-homologous end joining generates *MYC* rearrangements. Rearrangements resulted in over-expression of key genes and ChIP-seq identified that *HK2*, a member of the glucose metabolism pathway, is directly over-expressed through binding of *MYC* at its promoter.

## Introduction

The genome of multiple myeloma (MM) is characterized by primary translocations in ~40% of newly diagnosed patients that are considered initiating events and involve rearrangements of the immunoglobulin heavy chain (*IGH*) locus on 14q32.^1^ The partners of these rearrangements include 11q (*CCND1*, 15%), 4p (*FGFR3* and *MMSET*, 10%), 16q (*MAF*, 2-3%), 20q (*MAFB*, 1%) and 6q (*CCND3*, 1%). These rearrangements result in placement of the *IGH* super-enhancers next to a partner oncogene, resulting in its over-expression.^2^ The rearrangements predominantly occur in the switch regions 5’ of the constant regions in the *IGH* locus, where a high concentration of activation-induced cytidine deaminase (AID) binding motifs are found. Normally, AID binds to the switch regions leading to class switch recombination, resulting in antibody isotype switching.^3^ However, abnormal breaks in the switch regions, resulting from AID activity, results in *IGH* translocations.^4^

Secondary translocations involving *MYC*, located on 8q24.21, also occur in MM and are associated with disease progression and increased expression of *MYC*.^5–8^ *MYC* encodes a transcriptional regulator and has been shown to be involved in proliferation, differentiation, protein synthesis, apoptosis, adhesion, DNA repair, chromosomal instability, angiogenesis and metastasis.^9–13^ Translocations and high expression of *MYC* are associated with poor outcome, especially in MM where it is a marker of aggressive disease.^5, 14^ *MYC* can be deregulated by a range of different mechanisms including chromosomal rearrangement^5, 6^, copy-number gain/amplification^15, 16^, protein stabilization^17^, via secondary messengers involved in *MYC* transcription^18^ or miRNAs such as *PVT1*.^19, 20^

The frequency of *MYC* rearrangements seen in newly diagnosed MM (NDMM) varies from 15-50% and is dependent on the method used to identify it.^5, 6, 21, 22^ The data is consistent with *MYC* rearrangements being rare in the asymptomatic stages such as MGUS and smoldering myeloma^21^, and increases as the disease progresses with a high incidence (>80%) in myeloma cell lines.^22–24^

*MYC* rearrangements are not only seen in MM but are also frequent in lymphomas where they have been studied extensively.^25, 26^ In Burkitt’s lymphoma and diffuse large B cell lymphoma t(8;14) rearrangements between *IGH* and *MYC* have also been shown to result from abnormal class switch recombination.^27^ The relevance of AID in these rearrangements is supported by data from IL-6 transgenic mice which also develop *MYC*/*IGH* rearrangements in B cells. Rearrangements, however, do not occur if the mice are also deficient in AID, indicating that class switch recombination via AID is key in generating these rearrangements.^4, 28^ In MM, while karyotypic abnormalities similar to that observed in Burkitt’s lymphoma are seen, variant structures can also be detected, suggesting that the mechanism of rearrangement in MM may not be identical to lymphoma.^29^ Indeed, *MYC* rearrangements are not predominantly considered primary translocations in MM, often developing at later stages in the disease^22^, whereas in lymphoma they are considered primary events.^27^

We and others have previously shown that *MYC* translocations result in the juxtaposition of immunoglobulin loci super-enhancers to *MYC* resulting in its over-expression.^6, 30^ However, the details of breakpoint locations, the presence of copy-number abnormalities and the chromatin landscape of the rearrangement have not been well-characterized. In the present study, we have analyzed a large dataset of 1267 NDMM patients to determine the genomic architecture of *MYC* rearrangements and their effect on the expression of this proto-oncogene.

## Methods

### Patient Samples and Next Generation Sequencing

Total of 1267 NDMM patients were included in this study after informed consent and the study was approved by the Institutional Review Board at the University of Arkansas for Medical Sciences. Plasma cell were isolated from bone marrow by magnetic-activated cell sorting using CD138^+^ marker, AutoMACS Pro (Miltenyi Biotec GmbH, Bergisch Gladbach, Germany) or Robosep (STEMCELL Technologies, Vancouver, Canada). DNA from peripheral blood was used as a control sample for each patient to exclude germline variants. Three paired-end read sequencing platforms were combined without overlapping patients, namely targeted sequencing, whole exome sequencing, and low depth, long insert whole genome sequencing (**Supplementary Methods**). Additional expression data were available through either gene expression microarrays (Affymetrix, Santa Clara, CA, USA) or RNA-sequencing. An overall summary of methods, number of patients and external datasets are shown in **Supplementary Figure 1**. Patients’ characteristics are summarized in **Supplementary Table 1** and *MYC* region capture is illustrated in **Supplementary Figure 2**.

### Patient Derived Xenografts

Patient derived xenografts were generated by passaging primary patient CD138+ selected cells through the previously described SCID-rab myeloma mouse model.^31^ Tumors were dissected from the mouse, and pieces dispersed into a single cell population using a Kontes disposable tissue grinder. Cells were filtered through a 70 μm sterile filter, washed twice in PBS, treated with red cell lysis buffer, washed twice more, and treated immediately with Annexin V coated magnetic beads (Miltenyi Biotec), resulting in a population of cells with a viability >95%, as checked by flow cytometry. Passaged cells underwent CD138+ selection before being processed for 10x Genomics whole genome sequencing, RNA-sequencing, and ChIP-seq.

### ChIP-seq

ChIP-seq was performed on the myeloma cell lines KMS11 and MM.1S as well as a PDX sample with a *MYC* rearrangement identified by whole genome sequencing. 1×10^7^ cells per mark were fixed in a 1% formaldehyde solution, followed by the addition of glycine to a final concentration of 0.125 M. Cells were washed and resuspended in PBS containing 0.5% lgepal with 1% PMSF, before being pelleted and frozen at −80 °C. ChIP-seq for the histone marks H3K4me1, H3K4me3, H3K9me3, H3K27me3, H3K27Ac, and H3K36me3 (Active Motif, Carlsbad, CA, USA), as well as the super-enhancer proteins BRD4 and MED1 (Bethyl, Montgomer, TX, USA), and the transcription factor *MYC* (Santa Cruz Biotechnology, Dallas, TX, USA) were performed by Active Motif. Controls without antibody input were performed to ensure data quality.

### Data Analysis

Data analysis was performed as described previously, with minor differences between sequencing modalities.^32^ For details see **Supplementary Methods**.

### Statistical Analysis

Basic statistical analysis was performed using GraphPad Prism 7.01 (GraphPad Software, San Diego, CA, USA), R 3.4.4 and/or RStudio 1.1.442. Fisher’s exact test, the Mann-Whitney U test, Spearman’s rank correlation and Log-Rank test with Benjamini-Hochberg adjustment were used for data analysis and P values ≤0.05 were considered statistically significant.

### Data Access

Sequencing data have been deposited in the European Genomic Archive under the accession numbers EGAS00001001147, EGAS00001002859, or at dbGAP under Accession phs000748.v5.p4.

## Results

### *MYC* Rearrangements Are Usually Present as Inter-Chromosomal Translocations, Co-Occur with Secondary Genetic Events and Are Associated with Shorter Survival in Non-Hyperdiploid Cases

We examined a set of 1267 NDMM patient samples that had undergone either whole genome sequencing, exome sequencing, or targeted sequencing, of which the latter two methods involved capture of 2.3 Mb and 4.5 Mb, respectively, surrounding the *MYC* locus. Structural abnormalities involving the region surrounding *MYC*, including translocations, inversions, tandem-duplications and deletions, were detected in 36.0% (456/1267) of NDMM samples. Of those 456, 56.6% (258/456) had only a translocation, and 30.0% (137/456) had only an intra-chromosomal rearrangement. In 13.4% (61/456), both translocation and intra-locus rearrangement were present. Non-synonymous *MYC* mutations were rarely detected (0.7%, 9/1264), **Supplementary Table 2**.

The frequency of 8q24 abnormalities was significantly increased across ISS stages (I – 28.6%, II – 37.5% and III – 41.6%, P<0.001), and were higher in the IMWG high-risk (34.6%) and standard-risk (28.1%) groups than in the low-risk group (23.6%, P<0.05). The association of 8q24 abnormalities with these negative prognostic factors may suggest a worse outcome of patients with 8q24 abnormalities, however analysis of this did not confirm the assumption in this dataset (**Supplementary Figure 4A**). Also, 8q24 abnormalities were associated with lower, rather than higher, NF-κB pathway activation (**Supplementary Figure 3**). Additional analysis, however, showed a significant effect of 8q24 abnormalities within the non-hyperdiploid sub-group (Figure 1A).

**Figure 1:**
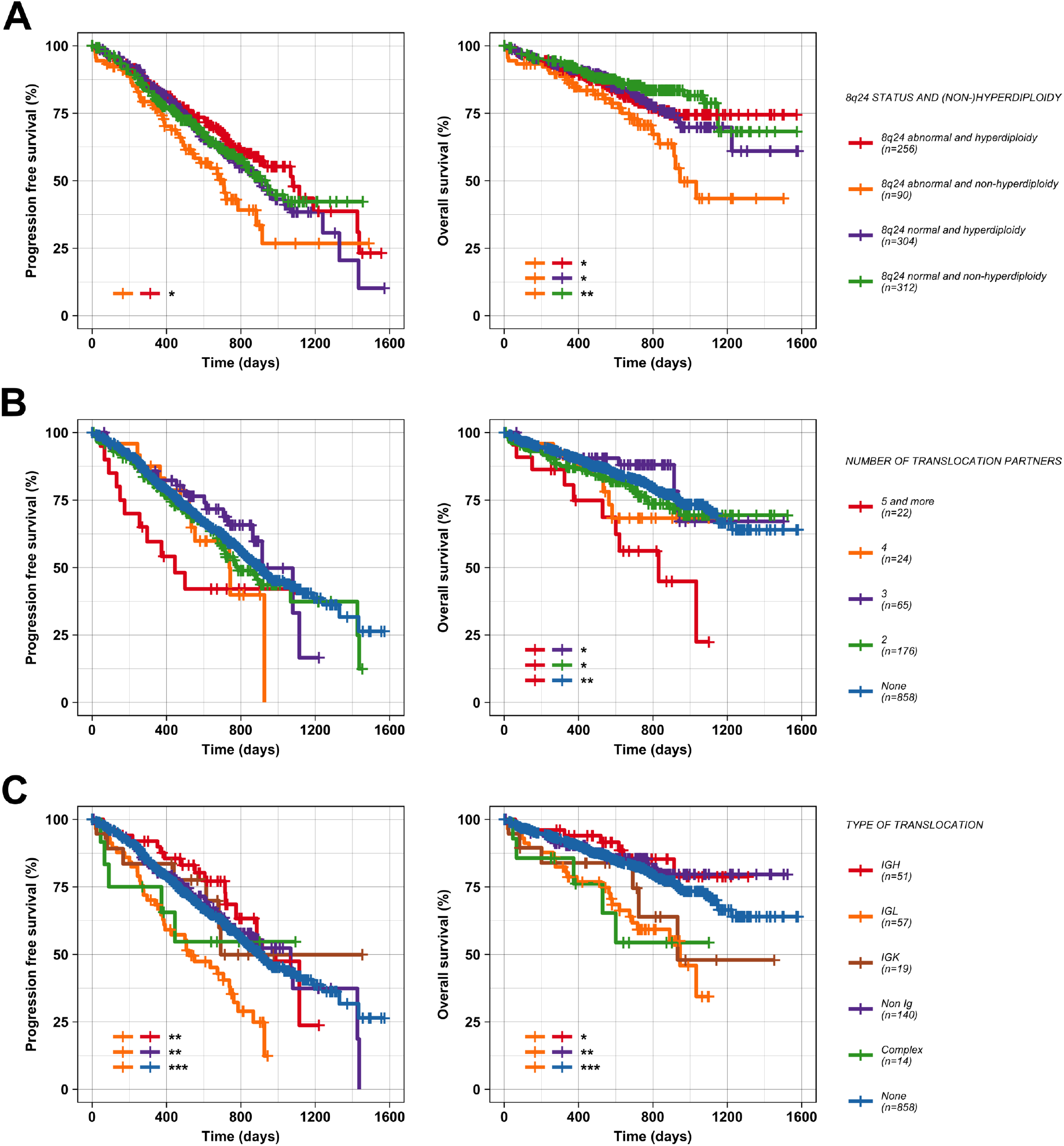
Effect of 8q24 abnormalities on patients’ outcome. **(A)** 8q24 abnormalities and hyperdiploidy. **(B)** Translocation complexity. **(C)** Translocations involving specific types of immunoglobulin locus. Statistically significant levels are as follows: ***P<0.001, **P<0.01 and *P<0.05.

Translocations were found in 25.2% (319/1267) of samples and occurred most frequently as inter-chromosomal translocations involving 2 to 5 chromosomes (90.3%, 288/319), but 4.4% (14/319) were highly complex and involved more than 5 chromosomal loci, Figure 2. Of the remaining cases, 5.3% (17/319) involved a large inversion of chromosome 8, >10 Mb in size. The proportion of *MYC* translocations involving 2, 3, 4, and 5 loci was 62.1% (198/319), 22.9% (73/319), 8.2% (26/319) and 2.5% (8/319), respectively. However, the number of chromosomes detected as affected by rearrangements involving *MYC* was dependent on the sequencing capture method used, as rearrangements involving 5 or more chromosomes were detected only by whole genome sequencing, **Supplementary Tables 3**–**4**. This data demonstrates that *MYC* is affected through chromoplexy, where 3 or more loci are involved in rearrangements, in 9.6% (121/1267) of NDMM or 26.5% (121/456) of samples with *MYC* abnormalities.

**Figure 2:**
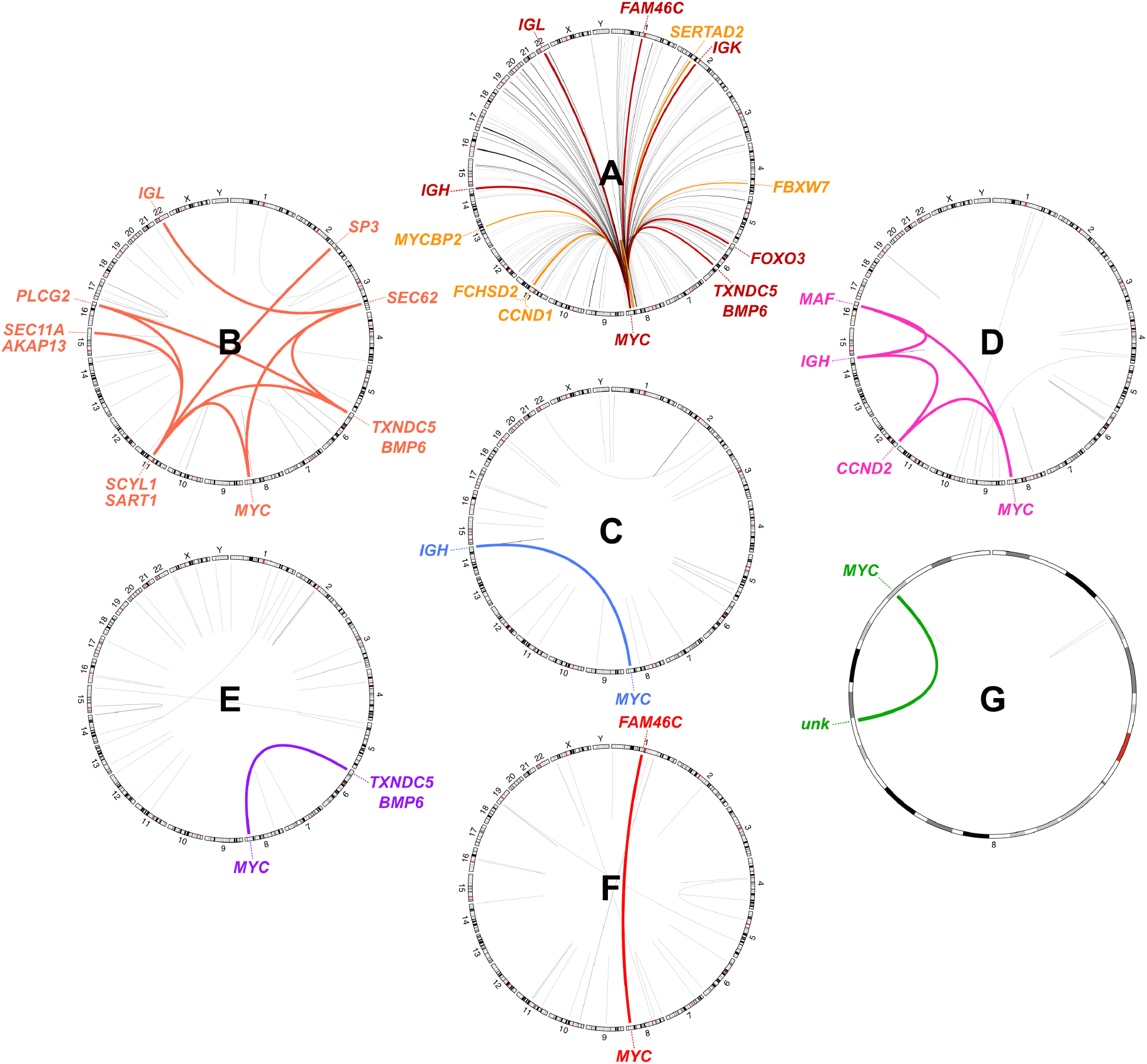
Circos plots of MM samples showing various *MYC* rearrangements. **(A)** *MYC* translocations partners in the dataset of the 1253 non-complex cases; loci present in 5– 9 cases (orange lines) and ≥10 cases (red lines) are highlighted. **(B)** Complex chromoplexy involving seven chromosomes, including the *MYC* locus. **(C)** Simple *IGH*-*MYC* t(8;14). **(D)** t(14;16) with a secondary translocation to *MYC*. **(E)** Non-Ig *MYC* translocation involving *TXNDC5*/*BMP6* on chromosome 6. **(F)** Non-Ig *MYC* translocation involving *FAM46C* on chromosome 1. **(G)** Inversion on chromosome 8. Annotated genes in uncertain loci were chosen as the closest highly-expressed gene(s) (within 1 Mb maximum distance) defined as being present in >95% of patients with log_2_ normalized counts >10 in the dataset of 571 cases tested by RNA-sequencing.

### *IGH-MYC* Translocation Breakpoints Have a Distinct Distribution Compared to Primary Translocations and Involve Recurrent Partners with Known Super-Enhancers

A total of 149 chromosomal loci were found to be involved in *MYC* translocations (Figure 2A; **Supplementary Table 5–6**). Six translocation partners were found in at least 10 cases and were the Immunoglobulin loci, *IGH* (63/1253, 5.0%), *IGL* (63/1253, 5.0%), *IGK* (26/1253, 2.1%), and also *TXNDC5/BMP6* on chromosome 6 (34/1253, 2.7%), *FAM46C* on chromosome 1 (20/1253, 1.6%), and *FOXO3* on chromosome 6 (14/1253, 1.1%), **Supplementary Table 5**. Each of these non-Ig loci were confirmed to contain highly-expressed genes in MM using RNA-sequencing data, being present in >95% of patients with log_2_ normalized counts >10. All of the loci except for *IGK* had super-enhancers previously identified in the MM.1S cell line. 67.2% (205/305) of cases with non-complex translocation (5 or less loci involved) had at least one of these super-enhancers involved in the translocation. Another five partners were present in 5–10 cases, three of which overlapped with the highly-expressed genes *FCHSD2*, *FBXW7* and *SERTAD2*, which are associated with known super-enhancers.^30^

Interestingly, 13 samples had complex *MYC* translocations with more than one of these super-enhancers. Also, eight samples had rearrangements involving *IGH*, *MYC* and *CCND1*, and four samples had rearrangements with *IGH*, *MYC* and *MAF*, indicating that they may occur as primary events early in the disease process. All oncogenes involved in these translocations show high expression (**Supplementary Figure 5**). This targeting of multiple oncogenes may explain worse patients’ survival with complex *MYC* translocations (Figure 1B). Ig loci were involved in 47.9% (146/305) of cases with a *MYC* translocation and were not associated with significantly higher *MYC* expression (Figure 3B, **Supplementary Figure 6B**) or patient s’ survival (**Supplementary Figure 4D**) compared to samples involving other super-enhancer-associated genes. In six cases, an *IGH* translocation occurred together with one of the light-chain immunoglobulin loci, but no sample involved both light chain loci. Within the Ig translocation groups, patients with *IGL* partners showed significantly worse outcome in comparison to *IGH* (P<0.05), other non-Ig translocations (P<0.01) and cases without *MYC* translocations (P<0.001, Figure 1C).

**Figure 3:**
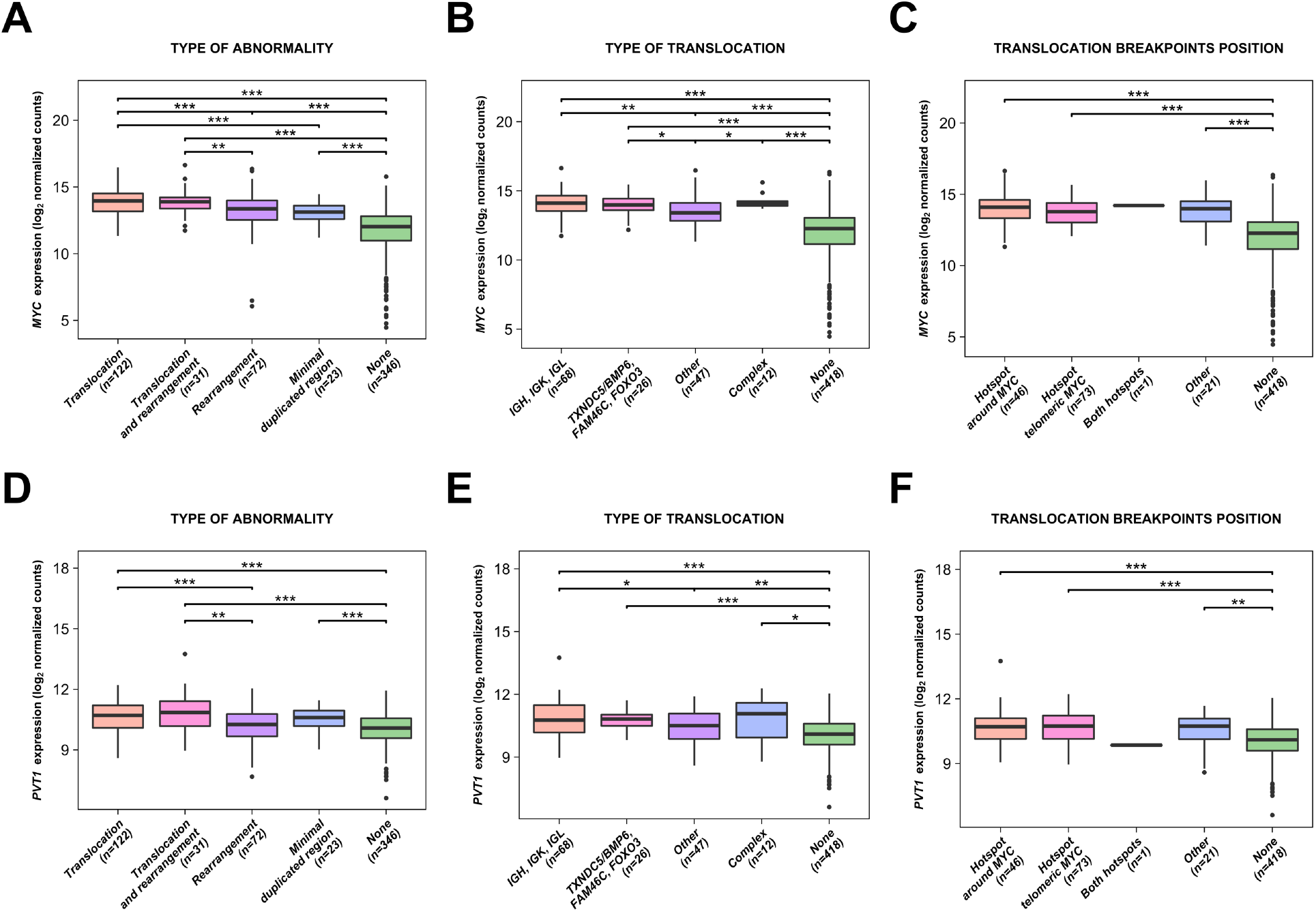
RNA-sequencing expression analysis of *MYC* and *PVT1* in relation to chromosomal abnormalities at 8q24. Effect of abnormality type [**(A)** and **(D)**], translocation category [**(B)** and **(E)**] and translocation breakpoint position [**(C)** and **(F)**] are shown for *MYC* and *PVT1*, respectively. Statistically significant levels are as follows: ***P<0.001, **P<0.01 and *P<0.05.

Analysis of the breakpoints at the *IGH* locus indicated a different pattern of *MYC* rearrangements to that of the primary Ig translocations. The primary translocations involving t(4;14), t(6;14), t(11;14), t(14;16) and t(14;20) have breakpoints clustered around the constant switch regions where AID motifs are concentrated. However, the *MYC* translocations do not share this pattern and are dispersed across the constant region, showing no association with AID motif clusters. This indicates that the *MYC* translocations are likely independent of AID and occur in a manner that is distinct to that of the primary translocations, Figure 4A–B.

**Figure 4:**
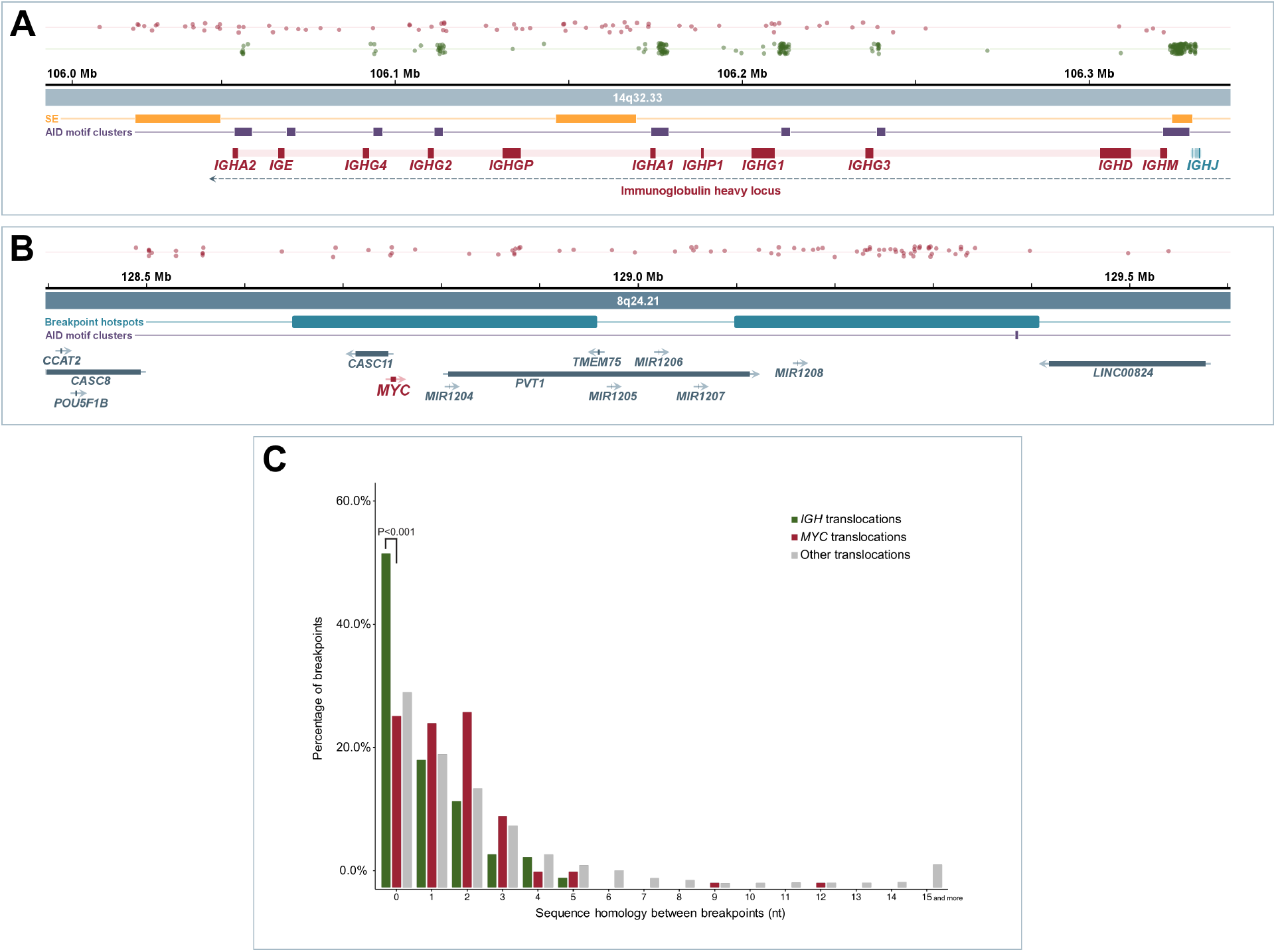
Primary *IGH* rearrangements and *MYC* rearrangements occur through different mechanisms. **(A)** The locations of classical *IGH* (green dots) and *IGH*-*MYC* (red dots) translocation breakpoints on 14q32.33. Yellow bars show super enhancers identified in MM.1S cell line; Purple bars show AID motif clusters (>200 RGYW motifs per 2.5 kb) indicating switch (S-) regions. *IGH* constant regions are indicated as red blocks. **(B)** *IGH*-*MYC* breakpoints on 8q24.21 (red dots). Blue bars show the two breakpoint hotspots identified in Figure 5. The location of *MYC* (red) and other genes (gray) are indicated. **(C)** Primary *IGH* translocations, *MYC* translocations and other translocations were compared for microhomology between chromosomes surrounding the breakpoints. Primary translocations have significantly more blunt-ended rearrangements compared to *MYC* rearrangements (P<0.001), consistent with MMEJ.

**Figure 5:**
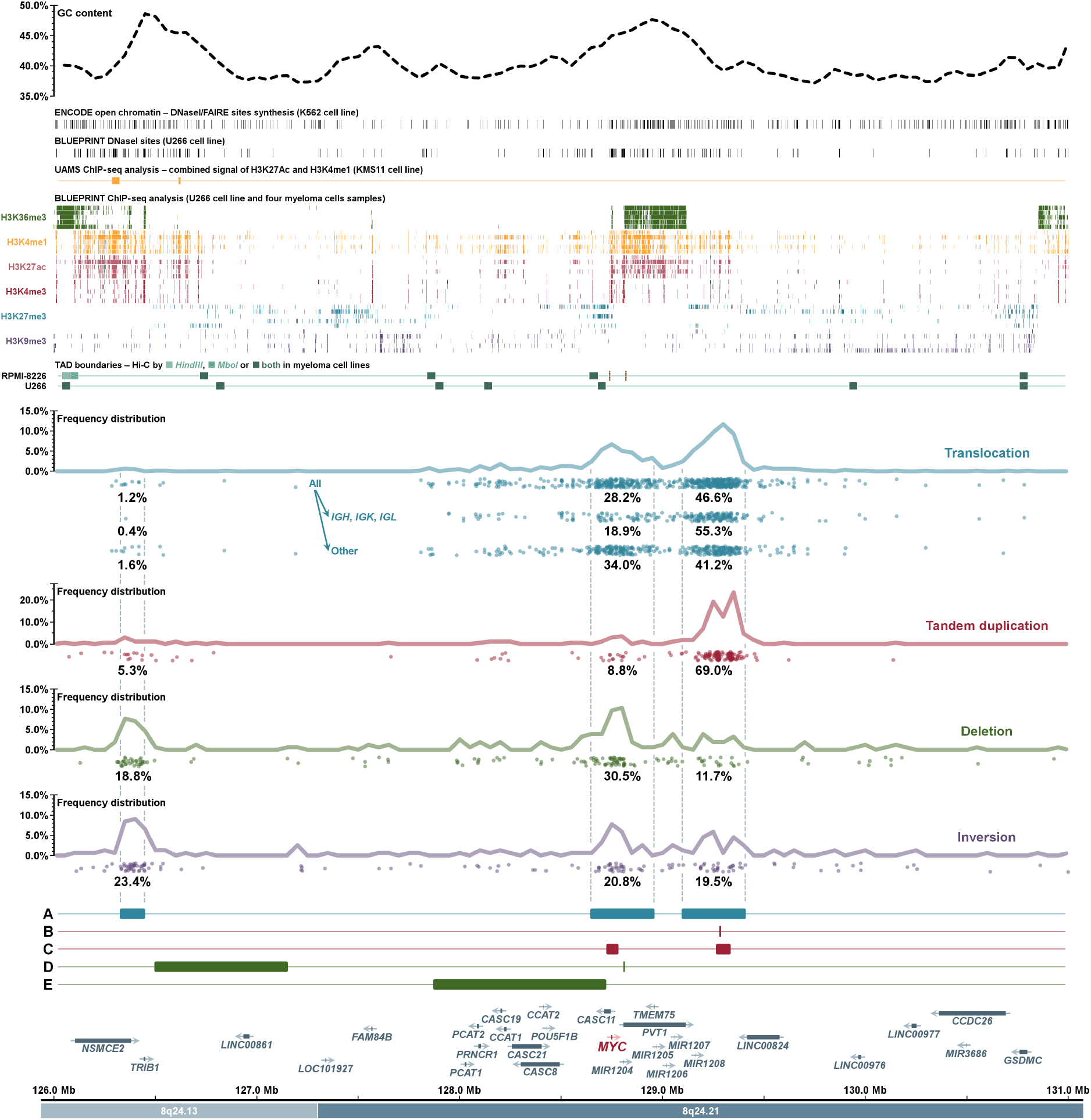
Distribution of chromosomal breakpoints and minimally altered regions detected at the *MYC* region. Percent values show proportion of breakpoints in the defined hotspot for a specific category of abnormalities. **(A)** Three breakpoints hotspots. **(B)** Minimal tandem-duplicated region. **(C)** Two minimal copy number gained regions (excluding tandem-duplications). **(D)** Two minimally deleted regions. **(E)** Minimal copy-number lost region (excluding deletions). Details of copy-number abnormalities analysis are given in Supplementary Figure 2–3. Upper dotted line shows GC content, ENCODE open chromatin markers identified by a combination of DNase-seq and FAIRE-seq in cell line K562, BLUEPRINT DNase-seq analysis of U266 cell line and BLUEPRINT ChIP-seq analysis in U266 cell line and four myeloma patients’ samples.

### *MYC* Breakpoints Show Evidence of Recombination through Microhomology

It is known that class switch recombination breakpoints in B cells occur through AID and non-homologous end joining (NHEJ), resulting in blunt ended DNA being ligated together.^33^ As the *MYC* breakpoints identified here do not align to switch regions, and are presumably not mediated by AID, we examined the aligned breakpoints to determine if they were constructed through blunt ended joining or other mechanisms. In comparison to re-aligned t(4;14), t(6;14), t(11;14), t(14;16) and t(14;20) breakpoints, which are mediated by AID and NHEJ, the *MYC* breakpoints had significantly fewer blunt ended rearrangements (54.1% vs. 27.7%, P<0.001) and significantly more rearrangements with at least two nucleotides of homology (25.4% vs. 45.8%) between the chromosomes, Figure 4C. Homologous sequences between chromosomes of up to 12 nts were found. Representative alignments of rearrangements are shown in the **Supplementary Alignments**. These homologous sequences are representative of microhomology-mediated end joining (MMEJ), which is a mechanism more common to all secondary translocation events, Figure 4C.

### 8q24 Breakpoints Occur in Three Hotspots and Associate with Open Chromatin Markers

Breakpoints were determined in a region covering up to 2.5 Mb from *MYC* and were categorized by the type of rearrangement. Three clusters of chromosomal breakpoints related to translocations, inversions, deletions and tandem-duplications were identified in the region chr8:126.0–131.0 Mb, Figure 5.

Translocation breakpoint hotspots were located in two 310 kb regions, one around *MYC* (chr8:128.6–129.0 Mb) and one telomeric of *MYC* (chr8:129.1–129.4 Mb). When examining all translocations, 28.2% were centered around the first hotspot and 46.6% around the second hotspot. However, there was an enrichment of Ig partner breakpoints at the second hotspot (55.3%) compared to first hotspot (18.9%), which was not so pronounced with non-Ig partners (41.2% vs. 34.0%). There was no evidence of an AID motif cluster at the second hotspot, which could have explained the enrichment for Ig partners and there was no effect of the breakpoint position on patient outcome (**Supplementary Figure 4E**).

Tandem-duplication breakpoints were enriched at the second hotspot (69.0% of breakpoints), Figure 5 and **Supplementary Figures 7–8**, as have been noted in MM cell lines previously.^34^ Conversely, deletion breakpoints were enriched at the first hotspot (30.5%) and at an additional hotspot centromeric of *MYC* (chr8:126.3–126.4 Mb). Inversion breakpoints were equally spread across all three hotspots.

By examining histone marks from the U266 cell line and four myeloma samples, for which we generated ChIP-seq histone mark data, there was also a link with accessible chromatin marks (H3K4me1, H3K4me3, H3K27ac and H3K36me3), DNaseI hypersensitivity sites and all three breakpoint hotspots, indicating that rearrangements may be more likely to happen in highly accessible, transcribed regions, Figure 5.

### Disruption of Topologically Associated Domains by *MYC* Rearrangements

Topologically associated domains (TADs) have been shown to contain DNA elements that are more likely to interact with one another. Disruption of these TADs may bring super-enhancer elements into the same TAD as *MYC*, resulting in its increased expression. We examined the super-enhancers from the MM.1S cell line and TADs from RPMI-8226 and U266 cell lines and integrated *MYC* breakpoints.

On the six frequent *MYC* translocation partner loci, breakpoints were clustered near to the super-enhancer and within the same TAD as the super-enhancer, Figure 6. At 8q24, the translocation breakpoints, at the two hotspots, were clustered within the TAD containing *MYC* and *PVT1*. The resulting rearrangements would bring the super-enhancer from the partner loci adjacent to *MYC*, resulting in the formation of a Neo-TAD (Figure 7B) and over-expression of *MYC*.

**Figure 6:**
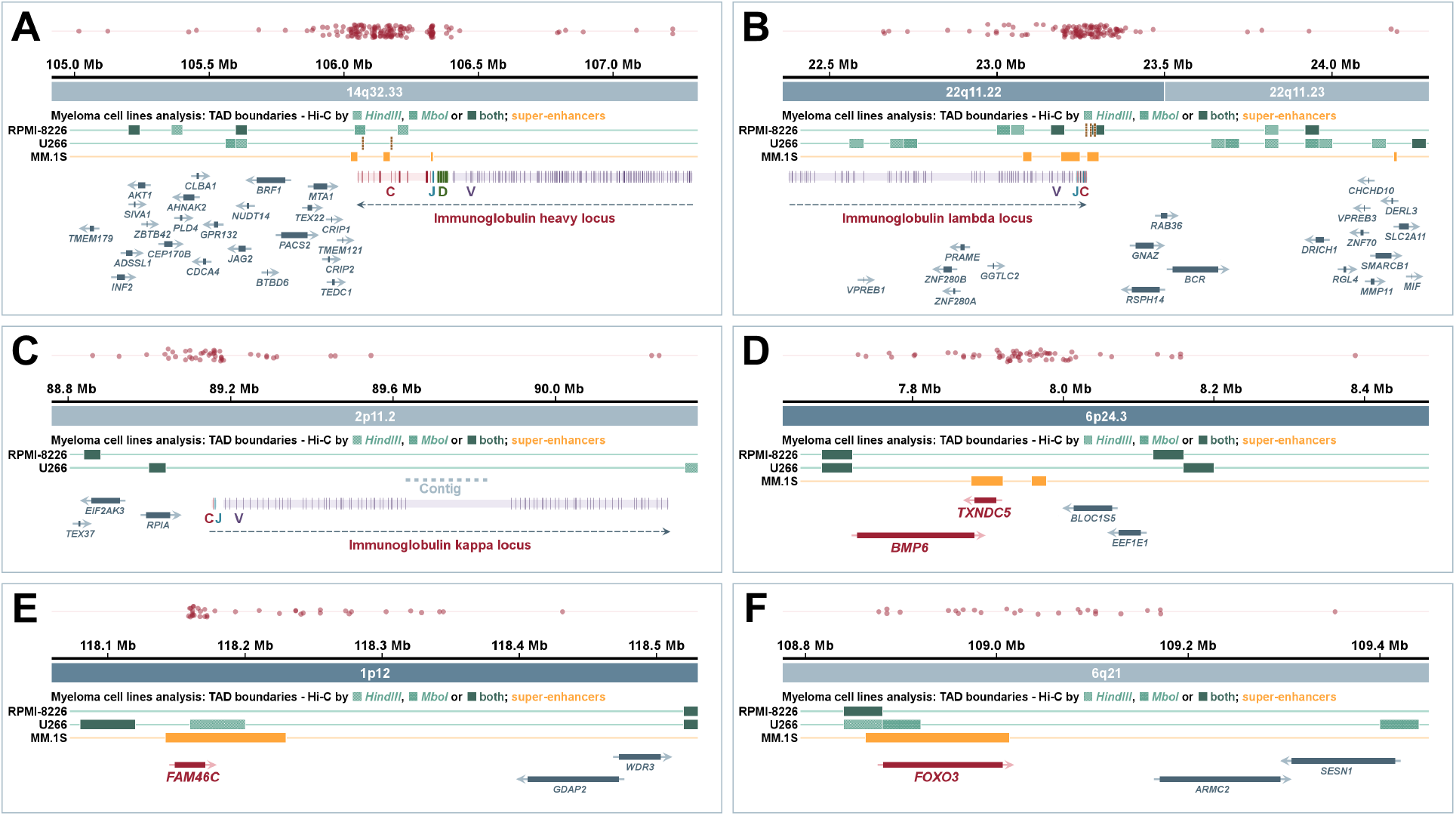
Chromosomal breakpoints in *MYC* translocation partners’ regions. **(A)** *IGH* locus at 14q32.33. **(B)** *IGL* locus on 22q11.22–22q11.23. **(C)** *IGK* locus on 2p11.2. **(D)** *TXNDC5*/*BMP6* locus on 6p24.3. **(E)** *FAM46C* locus on 1p12. **(F)** *FOXO3* locus on 6q21. Yellow bars show super-enhancers identified in the MM.1S cell line; Green bars show topologically associated domain (TAD) boundaries identified in RMPI-8226 and U266 cell lines. Ig genes are separated into constant (C, red), joining (J, blue), diversity (D, green) and variable (V, purple) regions; non-Ig highly-expressed genes (present in >95% of patients with log_2_ normalized counts >10 in the dataset of 571 cases tested by RNA-sequencing) are in red and other genes in gray color.

**Figure 7:**
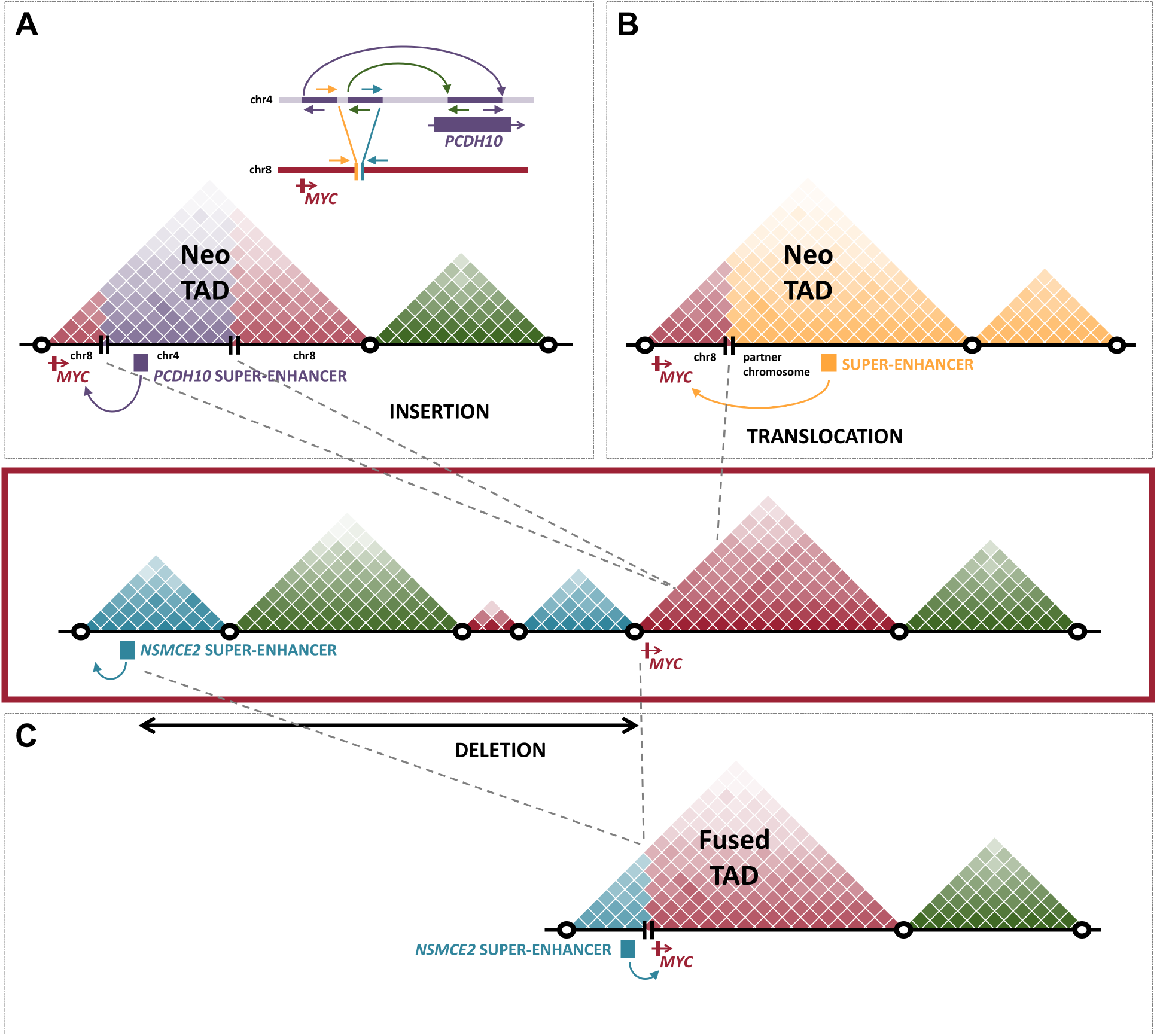
TAD reorganization through rearrangements places a super-enhancer next to *MYC*. The TAD architecture (colored triangles) surrounding *MYC* is indicated in the central panel (red box) as defined in U266 cells. **(A)** A patient sample with a t(4;8) involves the insertion of a super-enhancer from *PCDH10* (chr4) into chr8, creating a neo-TAD containing *MYC* and the super-enhancer. **(B)** A translocation from a key *MYC* partner introduces a super-enhancer into the *MYC* TAD. **(C)** Deletions centromeric of *MYC* result in fusion of TADs containing *MYC* and the super-enhancer next to *NSMCE2*.

We identified a patient derived xenograft sample with a t(4;8) that resulted in insertion of 3 regions of chromosome 4 next to *MYC*, Figure 7A. This resulted in the super-enhancer from *PCDH10*, defined by the presence of H3K27Ac and MED1 marks, being placed next to *MYC*, resulting in over-expression. This shows for the first time in a patient sample a rearrangement that confirms the importance of placing of a super-enhancer next to *MYC*.

Lastly, deletions at 8q24 centromeric of MYC are present in 2.9% (36/1249) of samples, Figure 5 and **Supplementary Figure 7–8**, most frequently result in contraction of the region bringing *NSMCE2* into close proximity of *MYC*, Figure 7C. This interstitial deletion results in TAD disruption bringing the super-enhancer at *NSMCE2*, present in the cell lines KMS11 and MM.1S, into the same TAD as *MYC*, resulting in a fused TAD and over-expression of *MYC*.

### 8q24 Translocations Result in Increased Expression of *MYC* and *PVT1*

The biological consequence of rearrangements at 8q24 is thought to be increased expression of *MYC*, so we examined the available CoMMpass study RNA-sequencing data, Figure 3, and a set of microarray data, **Supplementary Figure 6**, and categorized samples by type and location of breakpoints. In addition to *MYC*, we examined the expression of other genes in the regions, but only found significant increases in *MYC* and the non-coding RNA, *PVT1*, Figure 3A–F, which were associated with particular types of rearrangements. Expression level of these two genes showed a significant but weak correlation (r=0.4, P<0.001).

The six *MYC* partner loci present in >10 samples (*IGH*, *IGK*, *IGL*, *TXNDC5*/*BMP6*, *FOXO3* and *FAM46C*) had significantly higher expression of *MYC* (P<0.001) and *PVT1* (P<0.001) compared to those without rearrangements or less frequent partners, Figure 3B,E. Complex rearrangements involving more than five loci also resulted in higher expression of *MYC* (P<0.001) and *PVT1* (P=0.02) compared to those without rearrangements, at levels equivalent to the frequent translocation partners indicating a selection pressure on these six loci for increased *MYC* expression. There was no difference in expression between samples with breakpoints at the hotspot around *MYC* or telomeric of *MYC*, Figure 3C,F. Expression trends were not different in hyperdiploid (**Supplementary Figure 9**) and non-hyperdiploid (**Supplementary Figure 10**) subgroups, but a comparison between specific *MYC* abnormality groups shows that *MYC* and *PVT1* expression is higher in hyperdiploidy group (**Supplementary Figure 11**).

### Integration of *MYC* Binding Sites with Over-expressed Genes Identifies Proliferation Markers as Key Targets

We went on to determine if there is a gene expression signature associated with *MYC* abnormalities. We compared samples with and without any structural change at 8q24 and adjusted for hyperdiploidy status, as *MYC* abnormalities were present twice as often in samples with hyperdiploidy (46.0%, 290/630) as compared to non-hyperdiploid samples (22.7%, 102/449; P<0.001). A total of 121 genes (113 protein-coding and 8 non-coding RNA genes) were significantly de-regulated with a fold-change threshold of 1.8, of which 31.4% (38/121) were up-regulated and 68.6% (83/121) were down-regulated, Figure 8A. No significant pathway enrichment was detected by Gene Ontology Consortium^35^ using both PANTHER^36, 37^ and Reactome^38^ pathway analysis (for details of each gene see **Supplementary Table 7**).

**Figure 8:**
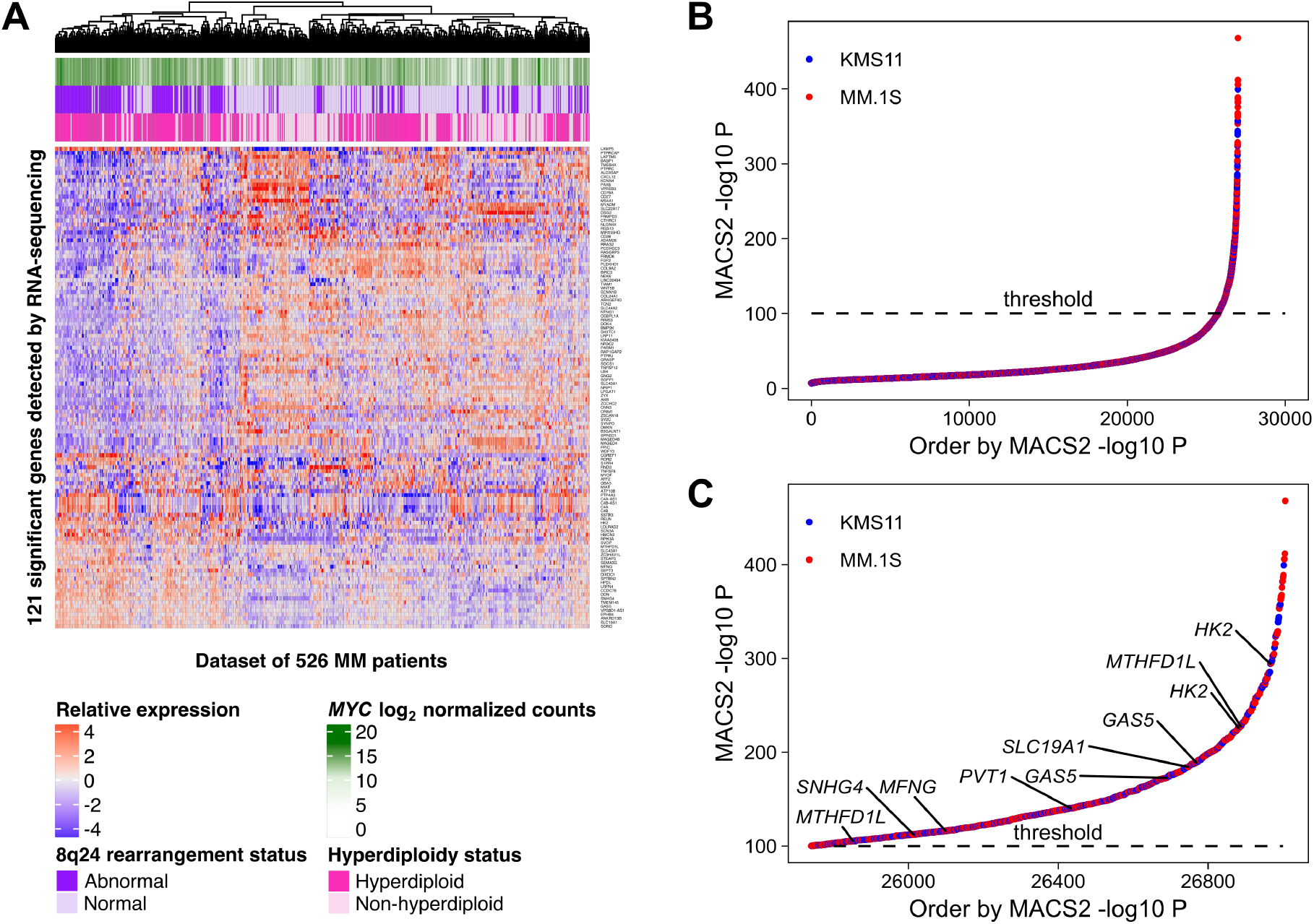
Integration of ChIP-seq for c-Myc and gene expression data identifies direct targets of *MYC* rearrangements. **(A)** 121 genes that were significantly changed in expression between samples with or without a *MYC* abnormality (FDR<0.05, fold-change ≥1.8) in the dataset of 526 MM patients with RNA-sequencing. **(B)** All c-Myc ChIP-seq peaks detected in MM.1S and KMS11 cell lines and ordered by −log10 P value. **(C)** Significant c-Myc ChIP-seq peaks (-log10 P value > 100) with highlighted *PVT1* gene and genes that overlap with 121 genes **(A)**.

We performed ChIP-seq against c-Myc and determined binding sites in two MM cell lines, MM.1S and KMS11, both of which have a *MYC* rearrangement. The peaks with a significance P < 10^−100^ using MACS2 in either cell line were considered significant and accounted for 4.7% of peaks (1266/27006), Figure 8B. The peaks were compared to the 121 genes that were significantly changed in expression, Figure 8A. Six genes were in the intersection between over-expressed and significant peaks: *HK2*, *MTHFD1L*, *SLC19A1*, *MFNG*, *SNHG4*, *GAS5*, Figure 8C. Using less stringent ≥1.3 fold-change cut-off that provided 1801 genes of which 40.8% (735/1801) were over-expressed, the intersection of over-expressed genes and those with a significant *MYC* binding peak was 25.3% (186/735). At the top of the list of 186 genes ordered by ChIP-seq −log10 P, we detected upregulation of the genes with known or potential oncogenic activity such as genes promoting cell proliferation, tumor growth and/or inhibition of apoptosis (*SNHG15*, *PPAN*, *MAT2A*, *METAP1D*, *MTHFD2*, *SNHG17*), translation factors (*EIF3B*, *EIF4A1*, *EEF1B2*) and genes involved in ribosome biosynthesis (*RPL10A*, *RPL35*, *RPL23A*, *RPSA*, *RPL13*, *WDR43*).

Importantly, we identified *HK2* and *PVT1* as direct targets of *MYC*. *HK2* is one of the most significant genes detected by ChIP-seq in both cell lines (-log10 P > 200, Figure 8C) as well as having the highest fold-change using RNA-sequencing analysis (**Supplementary Table 7**). This gene is an interesting direct target of *MYC* as it is part of the glucose metabolism pathway and would lead to increased energy metabolism and proliferation. *PVT1* showed a smaller fold-change by RNA-sequencing analysis (~1.4) but had a significant c-Myc protein binding site identified by ChIP-seq meaning that over-expression of *PVT1* is likely a downstream effect of *MYC* over-expression. This leads to a positive feedback loop and even higher *MYC* expression as *PVT1* positively regulates *MYC* expression.^39^

## Discussion

We show that *MYC* breakpoints in myeloma are clustered in three main hotspots on chromosome 8, one of which is associated with Ig translocations and tandem-duplications, another with non-Ig translocations and deletions, and the third with deletions and inversions. All breakpoints surrounding *MYC* result in increased expression of the oncogene, but inter-chromosomal translocations result in the largest increase in expression.

In this dataset we have used 1267 NDMM patient samples, of which 36.0% had *MYC* abnormalities, using next generation sequencing consisting of whole genome, exome and targeted panel data. The frequency of *MYC* abnormalities reported here is higher than previously seen using other techniques such as karyotyping or fluorescence *in situ* hybridization (FISH). This is likely due to the increased resolution of sequencing technologies, which can identify small insertions or deletions as well as translocations involving infrequent partner chromosomes. In addition, the complexity of breakpoints at 8q24 makes the placement of FISH probes difficult if all abnormalities are to be detected.

The scale of this analysis has allowed us to define the molecular breakpoints surrounding *MYC* with unparalleled accuracy and without technical bias. One of the two rearrangement hotspots involved in inter-chromosomal translocations in MM is also seen in other B cell malignancies. In Burkitt’s lymphoma, two breakpoint clusters within exon 1 and intron 1 of *MYC* were defined, which corresponds in location to the non-Ig rearrangement hotspot in MM.^26^ The same cluster is seen in diffuse large B cell lymphoma, where other random breakpoints are also seen scattered both centromeric and telomeric of *MYC*.^25^ Both of these studies looked at relatively small numbers of samples (78 and 17, respectively) and used older techniques, such as long distance PCR and FISH, to detect the breakpoints. It may be that in other B cell malignancies there are also other breakpoint hotspots similar to MM.

The main chromosomal partner to *MYC* through inter-chromosomal rearrangements is chromosome 14, specifically the *IGH* locus. In Burkitt’s lymphoma the *IGH-MYC* breakpoints on this chromosome lie almost exclusively within the switch regions (87%), upstream of the *IGH* constant regions.^26^ The remaining 13% are within the joining region of the locus. These breakpoints are consistent with the *IGH-MYC* rearrangement being a primary event in Burkitt’s lymphoma, occurring in 70-80% of patients.^40^ In contrast, in MM we clearly see that *IGH-MYC* breakpoints within the *IGH* locus are not in the switch or joining regions. Instead, they are spread out across the constant regions of the locus. This spread is distinct from the five common primary translocation breakpoints in MM [t(4;14), t(11;14), etc.] which are restricted to the switch and joining regions. Even those with *MYC* breakpoints within switch regions (6.9% of *IGH-MYC* rearrangements) also have primary rearrangements or are hyperdiploid. This indicates that the *IGH-MYC* rearrangements are secondary events in MM and probably occur through a different molecular mechanism to the primary translocation events. It is known that the primary translocations in MM, and the *IGH-MYC* primary events in Burkitt’s lymphoma, are mediated by AID and class switch recombination.^2, 4, 41^ Therefore, the *IGH-MYC* rearrangements may occur through an as yet unknown, AID-independent, mechanism.

The mechanism driving *MYC* rearrangements is likely not to involve NHEJ, which would result in blunt ended rearrangements.^33^ We have shown that *MYC* rearrangements are more likely to have short homologous sequences in common to both partner chromosomes, which is not seen as frequently in the primary *IGH* translocations. Short homologous sequences are indicative of MMEJ^42^, rather than NHEJ, and results through fork stalling and template switching during DNA replication or through microhomology-mediated break induced repair.^43, 44^ The proteins involved in MMEJ include PARP1, Rad50, and Ercc1 whereas MMEJ is inhibited by functional *ATM*, *H2AX*, *53BP1* and *BRCA1*.^42^ We have previously shown that mutation of *ATM*, *BRCA1* and other genes involved in DNA homologous recombination are associated with in increased levels of loss of heterozygosity in MM patients.^45^ It is likely that disruption of this pathway is key in genomic instability and progression of disease.

The non-Ig chromosomal partners of *MYC* are not random and are known to contain super-enhancer elements.^5, 6^ From our analysis of the breakpoints at the most frequent non-Ig locations (6p24.3 (*TXNDC5*/*BMP6*), 1p12 (*FAM46C*), 6q21 (*FOXO3*)) we show that the breakpoints at these genes are also clustered. The breakpoints are, in general, contained within TADs which are more likely to interact with one another.^46, 47^ Each TAD at the partner chromosome contains a super-enhancer and breakpoints rarely fall outside of the TAD. The rearrangements are predicted to result in a changed TAD structure that places *MYC* in the same domain as the super-enhancer from the partner locus. If breakpoints were to occur outside of the TAD with the super-enhancer there would be a lower likelihood of it interacting with *MYC* and expression would not be enhanced.

We identified 149 partner loci for *MYC* rearrangements, but 67.2% of the samples with translocations involve one of the six main partners. The Ig partners have strong super-enhancers in MM, but there are many other active super-enhancers and so it is likely that these six main partners are constrained by chromatin structure. The breakpoints at 8q24 surround an epigenetically active region, defined by the active chromatin marks H3K27Ac, H3K36me3 and, H3K4me1 as well as DNaseI hypersensitivity sites. It may be that epigenetically active, and therefore accessible, loci are preferred translocation partners^48, 49^, and the nuclear localization of chromosomes may play a part too.^50^

Each of these different rearrangements results in over-expression of *MYC*. *MYC* is not the only gene at 8q24, and indeed *PVT1* is significantly over-expressed in our dataset. *PVT1* is a long non-coding RNA and is associated with inhibition of apoptosis and increased proliferation.^51^ It has also been shown that *PVT1* interacts with *MYC*, resulting in a stable protein, and that ablation of *PVT1* results in diminished tumorigenicity.^52^ It may be that the gene complex encompassing *MYC* and *PVT1* is required for oncogenesis and merits further study.

Other than *PVT1*, we also identified other genes that are direct targets of c-Myc and are over-expressed in 8q24-rearranged samples. These included *HK2*, a key enzyme involved in glucose metabolism. It has previously been shown that silencing of *HK2* sensitizes cancer cells to other drugs, and so over-expression of *HK2* in *MYC*-rearranged myeloma may be a key drug resistance mechanism.^53^ Additional genes involved in important cellular functions that increase the oncogenic potential of myeloma cells were also identified, such as ribosome biosynthesis and translation initiation are likely to contribute to the poor prognosis seen in *MYC*-rearranged myeloma.^5, 14^ Targeting *MYC* could therefore be and effective way to disrupt many essential tumor features in one hit.

This study provides evidence of complex chromosomal rearrangements at 8q24 as a key cause of *MYC* oncogenic up-regulation. Although we found that several *MYC* abnormalities are associated with prognosis in this dataset, including *MYC-IGL* and complex translocations, we have previously shown that the association is not independent of other genomic and clinical markers.^54^ However, it may be possible that with longer follow-up *MYC* abnormalities may be independently associated with overall survival and be a marker of poor outcome. We also show a specific pattern of chromosomal breakpoints suggesting the role of the chromatin landscape in tumorigenesis. The mechanism of DNA breaks is clearly different between *MYC* rearrangements, resulting from MMEJ rather than NHEJ, and differs in myeloma compared to primary *MYC* translocations in lymphoma.

## Acknowledgements

Funding support for the CoMMpass dataset was provided by the Myeloma Genome Project. The CoMMpass dataset was generated by the Multiple Myeloma Research Foundation in collaboration with the Multiple Myeloma Research Consortium.

## Author Contributions

AM, CA, FED, GJM and BAW conceived and designed the study; AM, CA PQ, AR, AH and AT collected clinical data, sequencing data and performed statistical testing; JK, DA and AT were responsible for collection and availability of CoMMpass study related data; JAD, KRR, SY and FED developed and maintained mouse models; RGT, SD and ET processed the samples and performed laboratory protocols; AM, CA, ET, JK, DA, GHJ, GJM and BAW analyzed and interpreted data; CA, MAB, CPW, KM and MT performed the bioinformatics of the sequencing data; All authors discussed the results, wrote, reviewed and approved the manuscript.

## References

1. Morgan GJ, Walker BA, Davies FE. The genetic architecture of multiple myeloma. Nat Rev Cancer. 2012;12(5):335–348.

2. Bergsagel PL, Chesi M, Nardini E, Brents LA, Kirby SL, Kuehl WM. Promiscuous translocations into immunoglobulin heavy chain switch regions in multiple myeloma. Proc Natl Acad Sci U S A. 1996;93(24):13931–13936.

3. Stavnezer J, Guikema JE, Schrader CE. Mechanism and regulation of class switch recombination. Annu Rev Immunol. 2008;26(261–292.

4. Ramiro AR, Jankovic M, Eisenreich T, et al. AID is required for c-myc/IgH chromosome translocations in vivo. Cell. 2004;118(4):431–438.

5. Walker BA, Wardell CP, Brioli A, et al. Translocations at 8q24 juxtapose MYC with genes that harbor superenhancers resulting in overexpression and poor prognosis in myeloma patients. Blood Cancer J. 2014;4(e191.

6. Affer M, Chesi M, Chen WG, et al. Promiscuous MYC locus rearrangements hijack enhancers but mostly super-enhancers to dysregulate MYC expression in multiple myeloma. Leukemia. 2014;28(8):1725–1735.

7. Chng WJ, Huang GF, Chung TH, et al. Clinical and biological implications of MYC activation: a common difference between MGUS and newly diagnosed multiple myeloma. Leukemia. 2011;25(6):1026–1035.

8. Kuehl WM, Brents LA, Chesi M, Huppi K, Bergsagel PL. Dysregulation of c-myc in multiple myeloma. CurrTopMicrobiolImmunol. 1997;224(277–282.

9. Yu Q, Ciemerych MA, Sicinski P. Ras and Myc can drive oncogenic cell proliferation through individual D-cyclins. Oncogene. 2005;24(47):7114–7119.

10. Nesbit CE, Tersak JM, Grove LE, Drzal A, Choi H, Prochownik EV. Genetic dissection of c-myc apoptotic pathways. Oncogene. 2000;19(28):3200–3212.

11. Baudino TA, McKay C, Pendeville-Samain H, et al. c-Myc is essential for vasculogenesis and angiogenesis during development and tumor progression. Genes Dev. 2002;16(19):2530–2543.

12. Karlsson A, Deb-Basu D, Cherry A, Turner S, Ford J, Felsher DW. Defective double-strand DNA break repair and chromosomal translocations by MYC overexpression. Proc Natl Acad Sci U S A. 2003;100(17):9974–9979.

13. Yin XY, Grove L, Datta NS, Long MW, Prochownik EV. C-myc overexpression and p53 loss cooperate to promote genomic instability. Oncogene. 1999;18(5):1177–1184.

14. Walker BA, Wardell CP, Murison A, et al. APOBEC family mutational signatures are associated with poor prognosis translocations in multiple myeloma. Nat Commun. 2015;6(6997.

15. Carrasco DR, Tonon G, Huang Y, et al. High-resolution genomic profiles define distinct clinico-pathogenetic subgroups of multiple myeloma patients. Cancer Cell. 2006;9(4):313–325.

16. Avet-Loiseau H, Li C, Magrangeas F, et al. Prognostic significance of copy-number alterations in multiple myeloma. J Clin Oncol. 2009;27(27):4585–4590.

17. Sears R, Nuckolls F, Haura E, Taya Y, Tamai K, Nevins JR. Multiple Ras-dependent phosphorylation pathways regulate Myc protein stability. Genes Dev. 2000;14(19):2501–2514.

18. Shaffer AL, Emre NC, Lamy L, et al. IRF4 addiction in multiple myeloma. Nature. 2008;454(7201):226–231.

19. Manier S, Powers JT, Sacco A, et al. The LIN28B/let-7 axis is a novel therapeutic pathway in multiple myeloma. Leukemia. 2017;31(4):853–860.

20. Segalla S, Pivetti S, Todoerti K, et al. The ribonuclease DIS3 promotes let-7 miRNA maturation by degrading the pluripotency factor LIN28B mRNA. Nucleic Acids Res. 2015;43(10):5182–5193.

21. Avet-Loiseau H, Gerson F, Magrangeas F, et al. Rearrangements of the c-myc oncogene are present in 15% of primary human multiple myeloma tumors. Blood. 2001;98(10):3082–3086.

22. Gabrea A, Martelli ML, Qi Y, et al. Secondary genomic rearrangements involving immunoglobulin or MYC loci show similar prevalences in hyperdiploid and nonhyperdiploid myeloma tumors. Genes Chromosomes Cancer. 2008;47(7):573–590.

23. Shou Y, Martelli ML, Gabrea A, et al. Diverse karyotypic abnormalities of the c-myc locus associated with c-myc dysregulation and tumor progression in multiple myeloma. Proc Natl Acad Sci U S A. 2000;97(1):228–233.

24. Dib A, Gabrea A, Glebov OK, Bergsagel PL, Kuehl WM. Characterization of MYC translocations in multiple myeloma cell lines. J Natl Cancer Inst Monogr. 2008;39):25–31.

25. Bertrand P, Bastard C, Maingonnat C, et al. Mapping of MYC breakpoints in 8q24 rearrangements involving non-immunoglobulin partners in B-cell lymphomas. Leukemia. 2007;21(3):515–523.

26. Busch K, Keller T, Fuchs U, et al. Identification of two distinct MYC breakpoint clusters and their association with various IGH breakpoint regions in the t(8;14) translocations in sporadic Burkitt-lymphoma. Leukemia. 2007;21(8):1739–1751.

27. Kuppers R, Dalla-Favera R. Mechanisms of chromosomal translocations in B cell lymphomas. Oncogene. 2001;20(40):5580–5594.

28. Robbiani DF, Nussenzweig MC. Chromosome translocation, B cell lymphoma, and activation-induced cytidine deaminase. Annu Rev Pathol. 2013;8(79–103.

29. Gabrea A, Leif Bergsagel P, Michael Kuehl W. Distinguishing primary and secondary translocations in multiple myeloma. DNA Repair (Amst). 2006;5(9-10):1225–1233.

30. Loven J, Hoke HA, Lin CY, et al. Selective inhibition of tumor oncogenes by disruption of superenhancers. Cell. 2013;153(2):320–334.

31. Yata K, Yaccoby S. The SCID-rab model: a novel in vivo system for primary human myeloma demonstrating growth of CD138-expressing malignant cells. Leukemia. 2004;18(11):1891–1897.

32. Walker BA, Mavrommatis K, Wardell CP, et al. Identification of novel mutational drivers reveals oncogene dependencies in multiple myeloma. Blood. 2018;

33. Yan CT, Boboila C, Souza EK, et al. IgH class switching and translocations use a robust non-classical end-joining pathway. Nature. 2007;449(7161):478–482.

34. Demchenko Y, Roschke A, Chen WD, Asmann Y, Bergsagel PL, Kuehl WM. Frequent occurrence of large duplications at reciprocal genomic rearrangement breakpoints in multiple myeloma and other tumors. Nucleic Acids Res. 2016;44(17):8189–8198.

35. Gene Ontology C. Gene Ontology Consortium: going forward. Nucleic Acids Res. 2015;43(Database issue):D1049–1056.

36. Mi H, Huang X, Muruganujan A, et al. PANTHER version 11: expanded annotation data from Gene Ontology and Reactome pathways, and data analysis tool enhancements. Nucleic Acids Res. 2017;45(D1):D183–D189.

37. Mi H, Thomas P. PANTHER pathway: an ontology-based pathway database coupled with data analysis tools. Methods Mol Biol. 2009;563(123–140.

38. Fabregat A, Sidiropoulos K, Garapati P, et al. The Reactome pathway Knowledgebase. Nucleic Acids Res. 2016;44(D1):D481–487.

39. Carramusa L, Contino F, Ferro A, et al. The PVT-1 oncogene is a Myc protein target that is overexpressed in transformed cells. J Cell Physiol. 2007;213(2):511–518.

40. Molyneux EM, Rochford R, Griffin B, et al. Burkitt’s lymphoma. Lancet. 2012;379(9822):1234–1244.

41. Kuehl WM, Bergsagel PL. Multiple myeloma: evolving genetic events and host interactions. NatRevCancer. 2002;2(3):175–187.

42. Ottaviani D, LeCain M, Sheer D. The role of microhomology in genomic structural variation. Trends Genet. 2014;30(3):85–94.

43. Lee JA, Carvalho CM, Lupski JR. A DNA replication mechanism for generating nonrecurrent rearrangements associated with genomic disorders. Cell. 2007;131(7):1235–1247.

44. Hastings PJ, Ira G, Lupski JR. A microhomology-mediated break-induced replication model for the origin of human copy number variation. PLoS Genet. 2009;5(1):e1000327.

45. Pawlyn C, Loehr A, Ashby C, et al. Loss of heterozygosity as a marker of homologous repair deficiency in multiple myeloma: a role for PARP inhibition? Leukemia. 2018;

46. Spielmann M, Lupianez DG, Mundlos S. Structural variation in the 3D genome. Nat Rev Genet. 2018;

47. Dixon JR, Selvaraj S, Yue F, et al. Topological domains in mammalian genomes identified by analysis of chromatin interactions. Nature. 2012;485(7398):376–380.

48. Daniel JA, Nussenzweig A. The AID-induced DNA damage response in chromatin. Mol Cell. 2013;50(3):309–321.

49. Lu Z, Lieber MR, Tsai AG, et al. Human lymphoid translocation fragile zones are hypomethylated and have accessible chromatin. Mol Cell Biol. 2015;35(7):1209–1222.

50. Martin LD, Harizanova J, Righolt CH, et al. Differential nuclear organization of translocation-prone genes in nonmalignant B cells from patients with t(14;16) as compared with t(4;14) or t(11;14) myeloma. Genes Chromosomes Cancer. 2013;52(6):523–537.

51. Guan Y, Kuo WL, Stilwell JL, et al. Amplification of PVT1 contributes to the pathophysiology of ovarian and breast cancer. Clin Cancer Res. 2007;13(19):5745–5755.

52. Tseng YY, Moriarity BS, Gong W, et al. PVT1 dependence in cancer with MYC copy-number increase. Nature. 2014;512(7512):82–86.

53. Peng Q, Zhou J, Zhou Q, Pan F, Zhong D, Liang H. Silencing hexokinase II gene sensitizes human colon cancer cells to 5-fluorouracil. Hepatogastroenterology. 2009;56(90):355–360.

54. Walker BA, Mavrommatis K, Wardell CP, et al. A high-risk, Double-Hit, group of newly diagnosed myeloma identified by genomic analysis. Leukemia. 2019;33(1):159–170.

