## Supplementary_Data for "Microhomology-mediated end joining drives complex rearrangements and over-expression of *MYC* and *PVT1* in multiple myeloma"

### Supplementary Methods

#### Patient Samples and Next Generation Sequencing

Total of 1267 NDMM were included in this study after informed consent. Plasma cell were isolated from bone marrow by magnetic-activated cell sorting using $CD138^{+}$ marker, AutoMACS Pro (Miltenyi Biotec GmbH, Bergisch Gladbach, Germany) or Robosep (STEMCELL Technologies, Vancouver, Canada). DNA from peripheral blood was used as a control sample for each patient to exclude germline variants. Three paired-end read sequencing platforms were combined without overlapping patients. Overall summary of methods, number of patients and external datasets are demonstrated **in Supplementary Figure 1**. Patients' characteristics are summarized in **Supplementary Table 1** and *MYC* region capture is illustrated in **Supplementary Figure 2**.

**a. Targeted sequencing (n=100):** DNA was isolated using AllPrep DNA/RNA Kit (Qiagen, Hilden, Germany). Total of 50 ng of DNA was enzymatic fragmented and library was prepared using KAPA HyperPlus Kit (Kapa Biosystems, Wilmington, MA, USA) and SeqCap EZ Kit (Roche NimbleGen, Basel, Switzerland). A total of 4.8 Mb was targeted and designed in two parts. First, 4.2 Mb covering *IGH*, *IGK*, *IGL* and *MYC* genes focusing on translocations and chromosomal structure abnormalities. Second, 0.6 Mb covering exonic regions of 127 MM-specific genes and 27 chromosome regions for gene mutations and copy-number abnormalities analysis. Hybridization reactions were performed separately for each targeted-enrichment part and samples were finally combined at appropriate ratio to get required depth for chromosome structure abnormalities (~100x) and gene mutations (~250x) part. HiSeq 2500 (Illumina, San Diego, CA, USA) was used for sequencing. The DNA quality and quantity were measured by Qubit Fluorometer (Thermo Fisher Scientific, Waltham, MA, USA) and/or 2200 Tapestation (Agilent Technologies, Santa Clara, CA, USA). With focus on *MYC*, 4.5 Mb region (chr8:126.3–130.8 Mb) surrounding the gene was targeted with 83.1% capture. *MYC* expression level was defined in 98 patients by gene expression profiling using U133Plus2.0 microarray platform (Affymetrix, Santa Clara, CA) as previously described.[1]

**b. Whole exome sequencing (n=461):** A previous published dataset of patients with custom-enriched exome sequencing was used with detailed description of the protocol.[2] Briefly, DNA was isolated using AllPrep DNA/RNA Kit (Qiagen, Hilden, Germany). A total of 200 ng of DNA was fragmented using Covaris E-Series. NEBNext DNA library prep master mix set for Illumina (New England Biolabs, Ipswich, MA, USA) was used for library preparation. Exome enrichment was performed by custom designed RNA baits (SureSelect Human All Exon V5, Agilent Technologies; enriched for *IGH*, *IGK*, *IGL* and *MYC* region capture). Samples were sequenced using a HiSeq 2000 (Illumina, San Diego, CA, USA). The DNA quality and quantity were measured by Pico-green (Thermo Fisher Scientific, Waltham, MA, USA) and/or 2200 Tapestation (Agilent Technologies, Santa Clara, CA, USA). A region 2.3 Mb (chr8:127.5–129.8 Mb) surrounding *MYC* with 100% capture was targeted.

**c. Genome sequencing (n=706):** Dataset of patients was provided by Multiple Myeloma Research Foundation CoMMpass study and it is composed of patients with varying treatment strategies including bortezomib or carfilzomib-based regimens that may have been combined with IMiDs. Long-insert-based genome sequencing data was used for *MYC* translocation and chromosomal abnormalities study of the region in size of 5.0 Mb surrounding *MYC* (chr8:126.0–131.0 Mb). Exome sequencing available in 703 of 706 patients was used for NS-SNVs analysis. Expression of genes was quantified by RNA-Sequencing available in 571 of 706 patients.

**Data Analysis**

Data analysis was performed as described previously, with minor differences between sequencing modalities.[3] Briefly, FASTQ files from targeted sequencing (TS), whole exome sequencing (WES) and whole genome sequencing (WGS) were aligned to the human genome assembly GRCh37 by BWA-MEM (v0.7.12). Variants were called using MuTect2 and Strelka (v1.0.14 in TS, v1.0.15 in WES and WGS), filtered using fpfilter (https://github.com/ckandoth/variant-filter) in TS and a custom filter described elsewhere in WES and WGS.[3] A minimum 10% VAF filter was used for indels. Variant annotation was provided by Variant Effect Predictor (v85) in TS or Oncotator (v1.9.0) in WES and WGS.

Intra- and inter-chromosomal rearrangements were called using Manta[4] (v0.29.6 in WES and v1.0.1 in WES and WGS) with default settings and the exome flag specified for TS and WES samples. Copy-number alterations were determined in TS by normalized tumor/germline depth ratio supported by allele ratio changes in individual heterozygous SNP loci. All *MYC*-region-associated chromosomal breakpoints and copy number abnormalities were manually inspected. Cases with more than five chromosomes involved in the translocation (n=14) or more than five intra-chromosomal rearrangements at 8q24 (n=18) were considered as abnormal, but for high inter- or intra-chromosomal complexity they were excluded from detailed analysis. *MYC* region annotations for the CoMMpass and UK datasets are detailed in a previous publication.[3]

Manta was used to evaluate sequence homology between breakpoints in WGS data. All passed translocation events were filtered to only include classic *IGH* or *MYC* translocations. All events with the IMPRECISE flag set in Manta were filtered out. The homology length (HOMLEN) parameter was extracted from the INFO field in the Manta VCF. Fields without a HOMLEN parameter were set to zero. To ensure viability of Manta homology detection we manually verified randomly selected samples (see **Supplementary Alignments**). Events with only one nucleotide homology between breakpoints were not considered for analysis due to the fact that those could be simply due to chance. Finally, *IGH* and *MYC* events with no sequence homology were compared to *IGH* and *MYC* events with two or more nucleotide homology using Fisher's exact test.

RNA-Sequencing data was aligned to the human genome assembly GRCh38 with gene-transcripts quantification processing by Star (v2.5.1b) and Salmon (v0.6.0) algorithms. The read counts per gene from Salmon were read into R and using the DESeq2 (v1.20.0) R library, normalized across samples and the $\log_2$ expression calculated. A total of 526 patients with available RNA-Sequencing data and hyperdiploidy status were analyzed for a *MYC* signature using limma R package. Genes with more than 0.5% of zero values were excluded from the analysis, remaining genes were adjusted for hyperdiploidy status and filtered by FDR≤0.05 and fold change >=1.8. Threshold $\log_2$=13.0 for *MYC*-expression-based signatures was discriminated by receiver operating characteristics (ROC) analysis (AUC=0.85) as intersection between the highest sensitivity (0.75) and specificity (0.82) to predict abnormal genomic profiles. Gene enrichment was performed by Gene Ontology Consortium analysis with Fisher's test with FDR multiple test correction (P≤0.05).

#### External Datasets

Genomic annotations of breakpoint regions were taken from previously published sources. Super-enhancer sites were taken from the MM.1S myeloma cell line.[5] TADs were taken from the MM cell lines U266 and RPMI-8226.[6] Chromatin marks were taken from the MM cell line U266 and four myeloma cell samples.[7, 8] Open chromatin was identified by a combination of DNase-Seq and FAIRE-Seq in cell line K562.[9]

#### References

1. Zhan F, Hardin J, Kordsmeier B, et al. Global gene expression profiling of multiple myeloma, monoclonal gammopathy of undetermined significance, and normal bone marrow plasma cells. Blood. 2002;99(5):1745-1757.
2. Walker BA, Boyle EM, Wardell CP, et al. Mutational Spectrum, Copy Number Changes, and Outcome: Results of a Sequencing Study of Patients With Newly Diagnosed Myeloma. J Clin Oncol. 2015;33(33):3911-3920.
3. Walker BA, Mavrommatis K, Wardell CP, et al. Identification of novel mutational drivers reveals oncogene dependencies in multiple myeloma. Blood. 2018;
4. Chen X, Schulz-Trieglaff O, Shaw R, et al. Manta: rapid detection of structural variants and indels for germline and cancer sequencing applications. Bioinformatics. 2016;32(8):1220-1222.
5. Loven J, Hoke HA, Lin CY, et al. Selective inhibition of tumor oncogenes by disruption of super-enhancers. Cell. 2013;153(2):320-334.
6. Wu P, Li T, Li R, et al. 3D genome of multiple myeloma reveals spatial genome disorganization associated with copy number variations. Nat Commun. 2017;8(1):1937.
7. Stunnenberg HG, International Human Epigenome C, Hirst M. The International Human Epigenome Consortium: A Blueprint for Scientific Collaboration and Discovery. Cell. 2016;167(7):1897.
8. Adams D, Altucci L, Antonarakis SE, et al. BLUEPRINT to decode the epigenetic signature written in blood. Nat Biotechnol. 2012;30(3):224-226.
9. Song L, Zhang Z, Grasfeder LL, et al. Open chromatin defined by DNaseI and FAIRE identifies regulatory elements that shape cell-type identity. Genome Res. 2011;21(10):1757-1767.

### Supplementary Figures

#### Supplementary Figure 1: Graphical overview of methods, internal and external datasets.

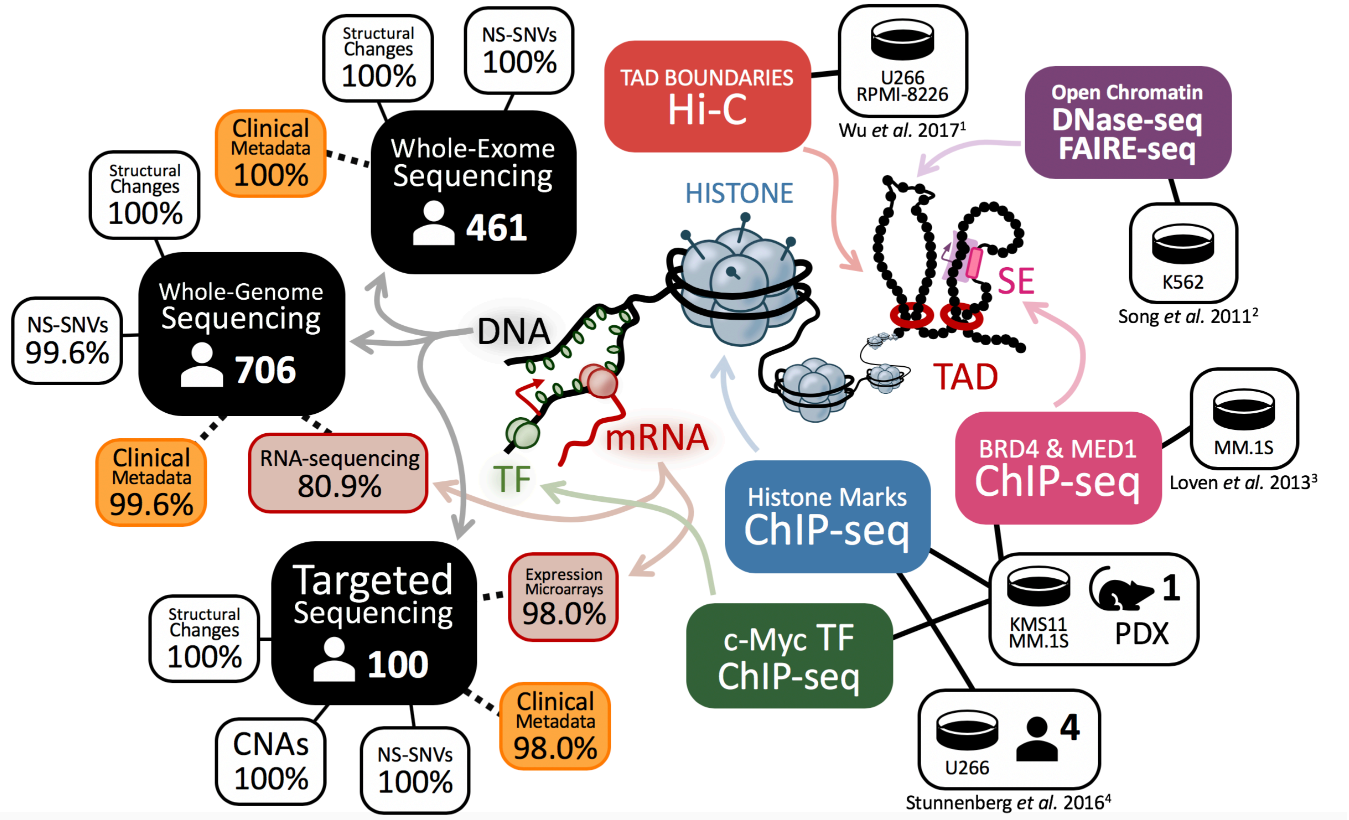

References:
1. Wu P, Li T, Li R, et al. 3D genome of multiple myeloma reveals spatial genome disorganization associated with copy number variations. Nat Commun. 2017;8(1):1937.
2. Song L, Zhang Z, Grasfeder LL, et al. Open chromatin defined by DNaseI and FAIRE identifies regulatory elements that shape cell-type identity. Genome Res. 2011;21(10):1757-1767.
3. Loven J, Hoke HA, Lin CY, et al. Selective inhibition of tumor oncogenes by disruption of super-enhancers. Cell. 2013;153(2):320-334.
4. Stunnenberg HG, International Human Epigenome C, Hirst M. The International Human Epigenome Consortium: A Blueprint for Scientific Collaboration and Discovery. Cell. 2016;167(7):1897.

**Supplementary Figure 2: Illustration of the studied *MYC* region at 8q24 in each dataset.** 461 cases with custom-enriched whole exome sequencing (up), 100 cases with targeted sequencing (middle), 706 cases with whole genome sequencing (down).

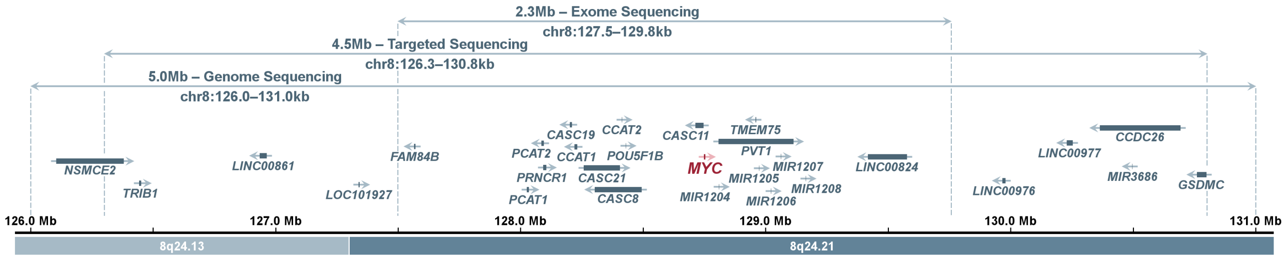

**Supplementary Figure 3: Association of 8q24 abnormalities and NF-κB pathway activation.** NF-κB pathways activation was defined as an average expression of the genes as follows: **(A)** NF-κB(11)[1] – *BIRC3*, *TNFAIP3*, *NFKB2*, *IL2RG*, *NFKBIE*, *RELB*, *NFKBIA*, *CD74*, *PLEK*, *MALT1*, *WNT10A*; **(B)** NF-κB(10)[2] – same as previous, excluding *BIRC3*; and **(C)** NF-κB(3)[2] – *TNFAIP3*, *IL2RG* and *BIRC3* (C). Expression was analyzed using RNA-sequencing. Statistically significant levels are as follows: ***P<0.001, **P<0.01 and *P<0.05.

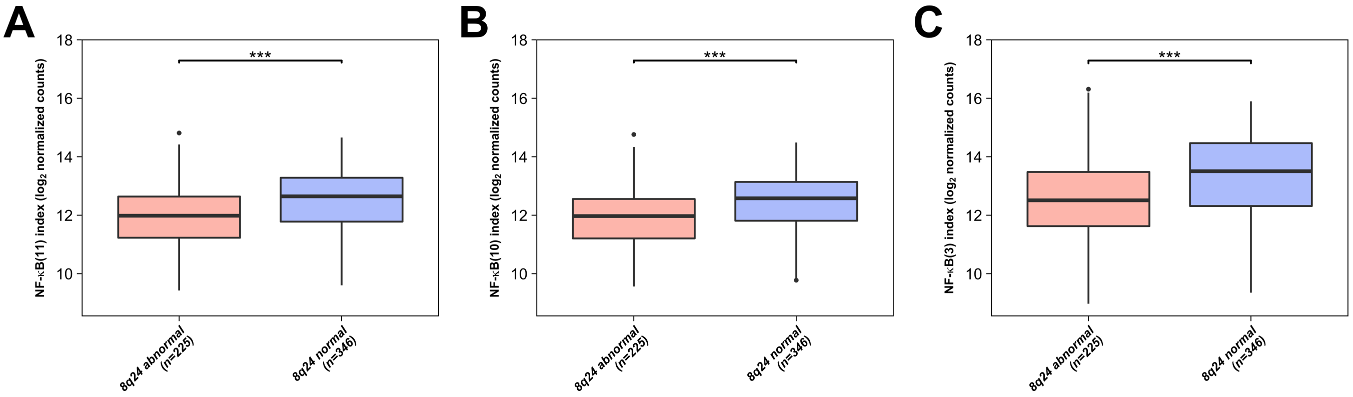

References:
1. Annunziata CM, Davis RE, Demchenko Y, et al. Frequent engagement of the classical and alternative NF-kappaB pathways by diverse genetic abnormalities in multiple myeloma. Cancer Cell. 2007;12(2):115-130.
2. Demchenko YN, Glebov OK, Zingone A, Keats JJ, Bergsagel PL, Kuehl WM. Classical and/or alternative NF-kappaB pathway activation in multiple myeloma. Blood. 2010;115(17):3541-3552.

**Supplementary Figure 4(A–C): Effect of 8q24 abnormalities on patients' outcome. (A)** 8q24 abnormalities. **(B)** Hyperdiploidy status. **(C)** Type of 8q24 abnormality. Statistically significant levels are as follows: ***P<0.001, **P<0.01 and *P<0.05. No significant P was found.

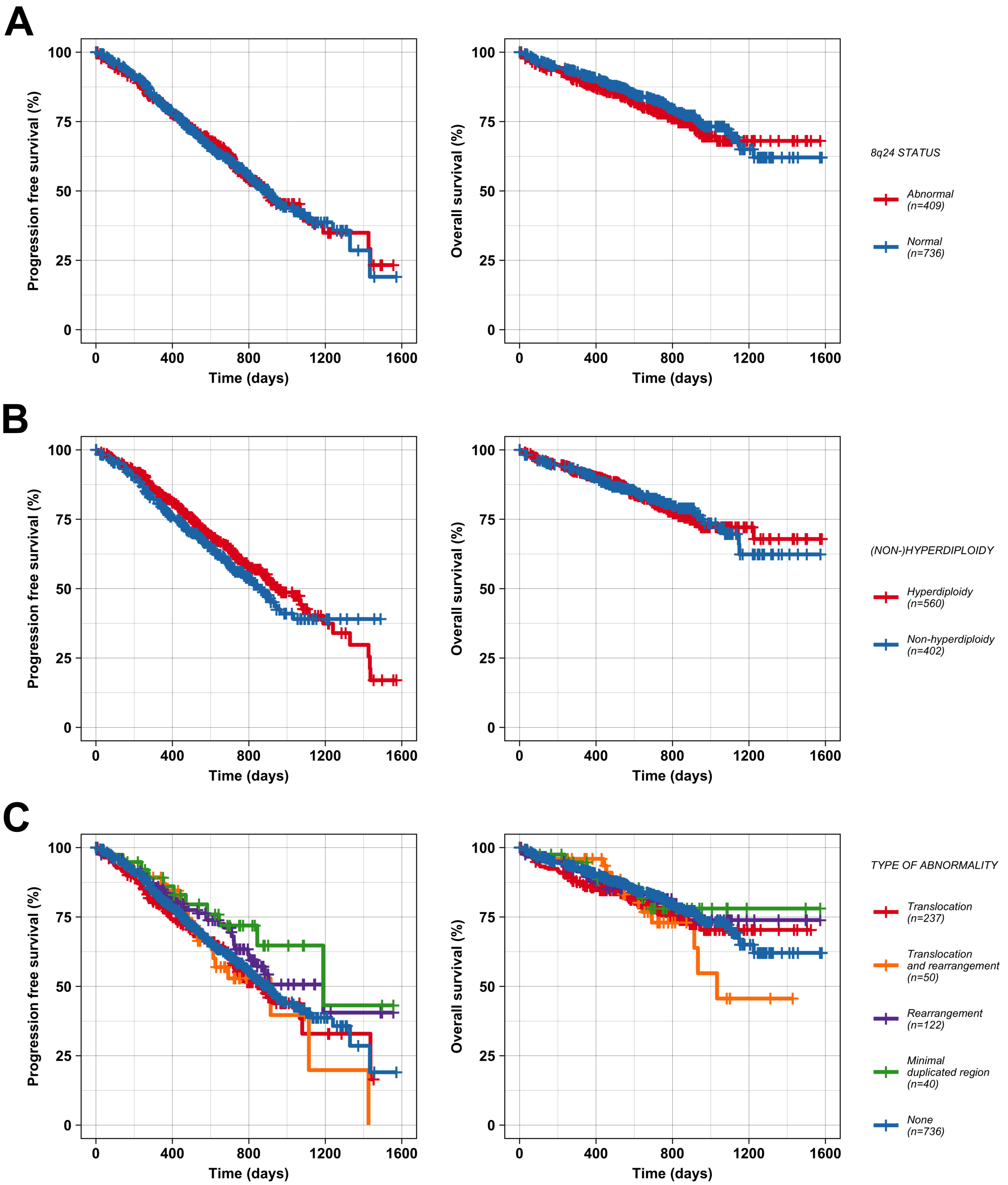

**Supplementary Figure 4(D–E): Effect of 8q24 abnormalities on patients' outcome. (D)** Translocation category. **(E)** Translocation breakpoint position. Statistically significant levels are as follows: ***P<0.001, **P<0.01 and *P<0.05. No significant P was found.

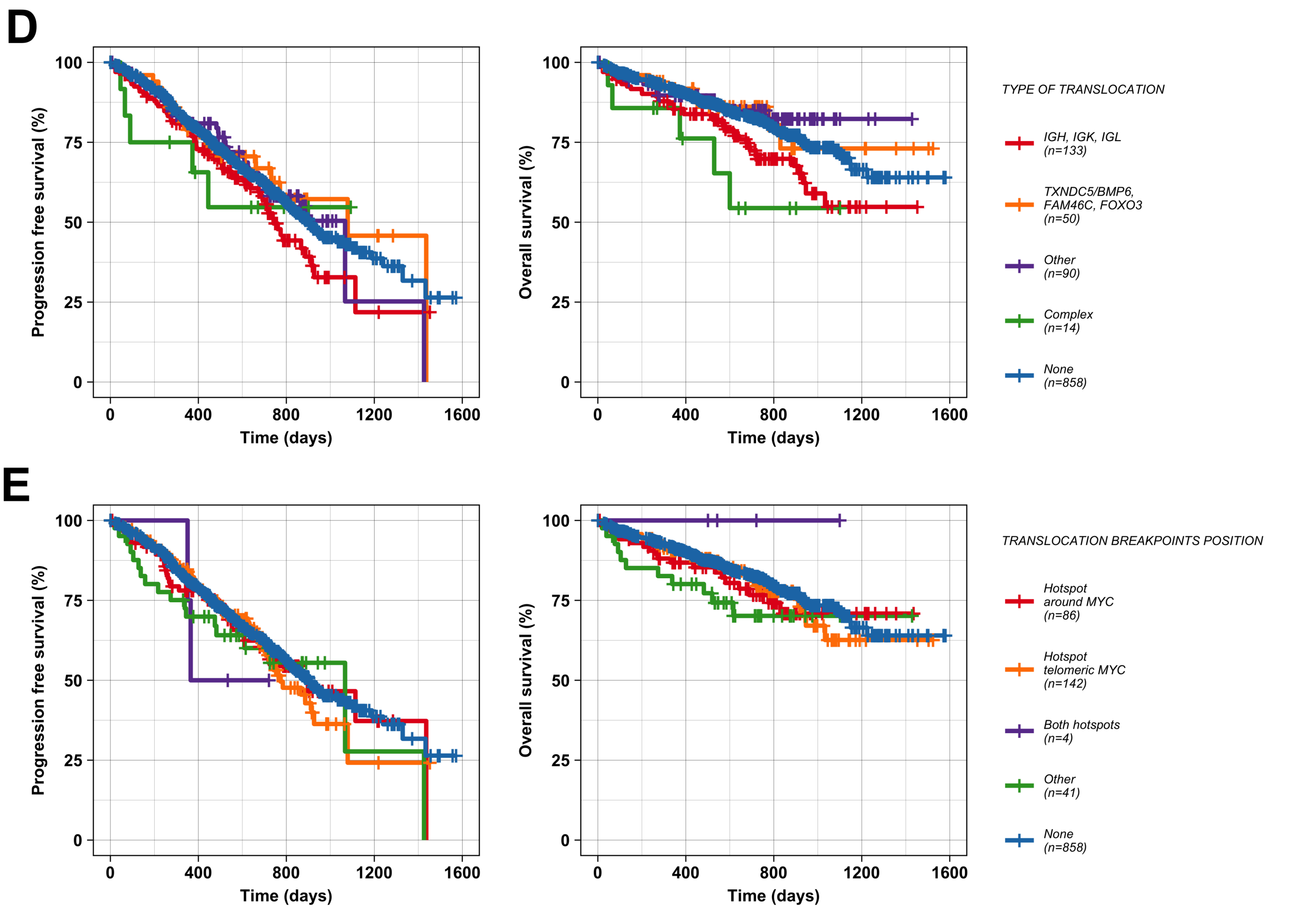

**Supplementary Figure 5(A–C): Expression of oncogenes in complex translocations in five cases with available RNA-sequencing data.** Box plots show expression distribution of the oncogene in specific *IGH* (left) and *MYC* (right) translocation groups. Red line determines a level of the oncogene expression in the case with complex translocation. Expression was analyzed using RNA-sequencing.

**Case A – t(8;14;16) *MYC-IGH-MAF***

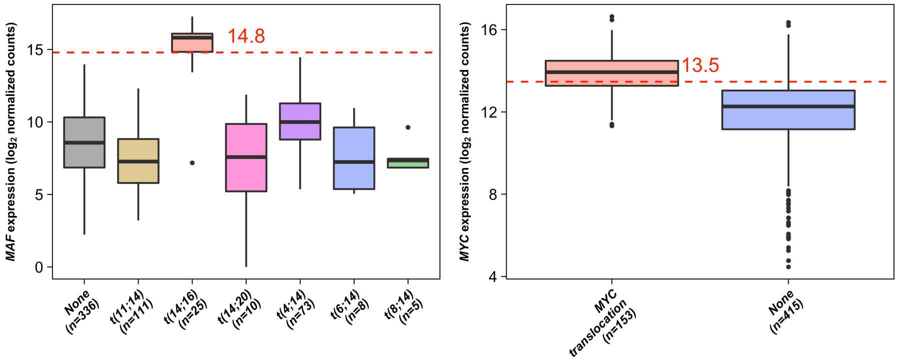

**Case B – t(8;14;16) *MYC-IGH-MAF***

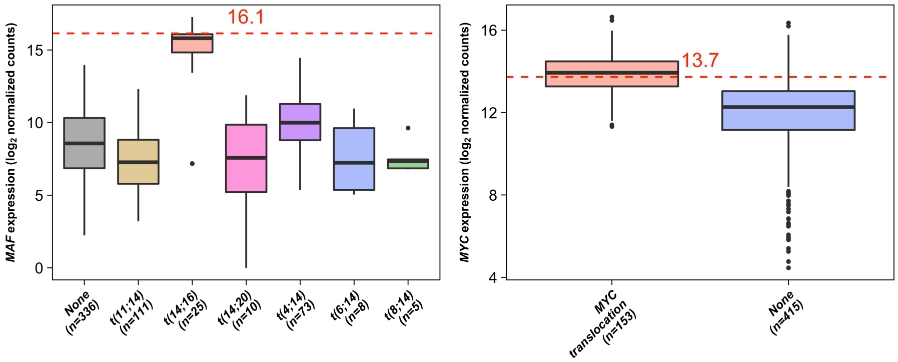

**Case C – t(8;11;14) *MYC-CCND1-IGH***

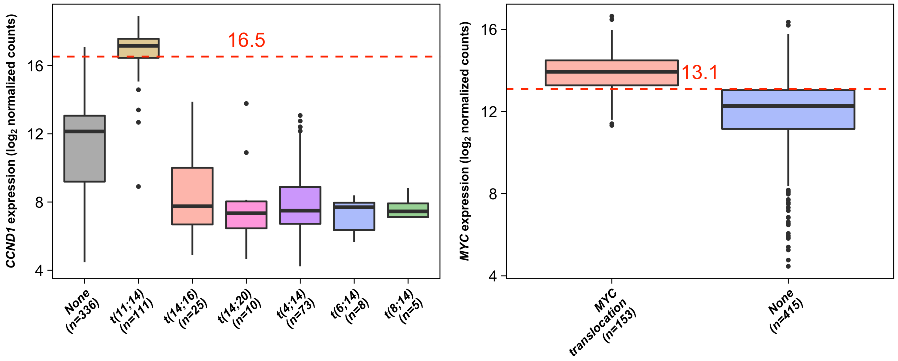

**Supplementary Figure 5(D–E): Expression of oncogenes in complex translocations in five cases with available RNA-sequencing data.** Box plots show expression distribution of the oncogene in specific *IGH* (left, middle) and *MYC* (right) translocation groups. Red line determines a level of the oncogene expression in the case with complex translocation. Expression was analyzed using RNA-sequencing.

**Case D – t(8;12;14;16) *MYC-CCND2-IGH-MAF***

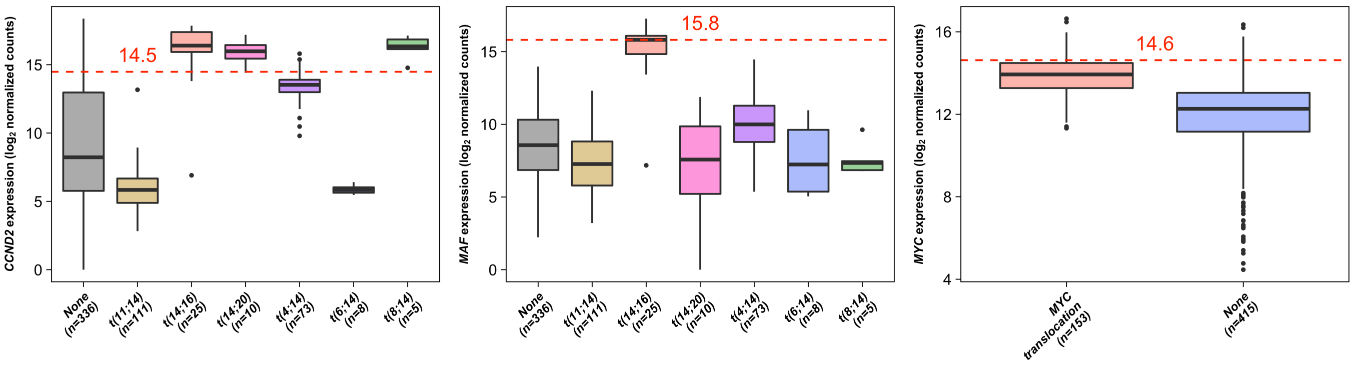

**Case E – t(8;11;14;18;22) *MYC-CCND1-IGH-BCL2-IGL***

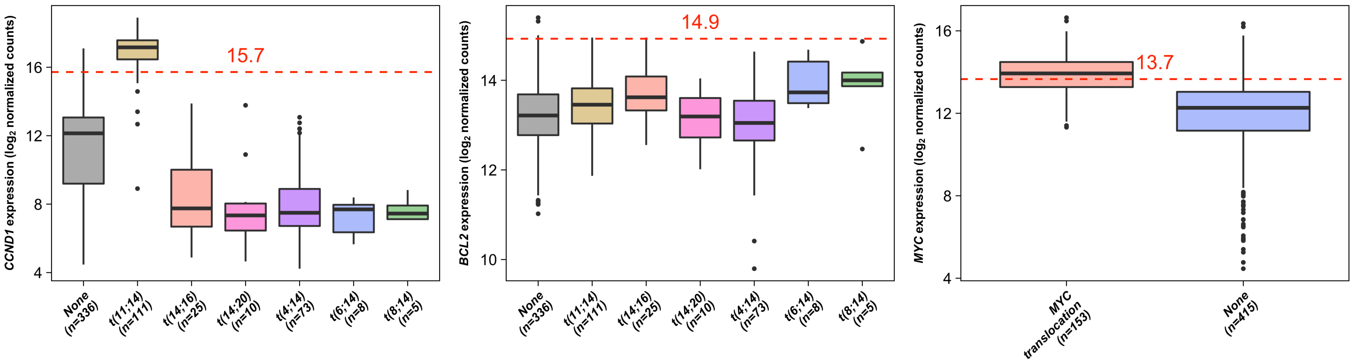

**Supplementary Figure 6: Gene-expression microarray analysis of *MYC* in relation to chromosomal abnormalities at 8q24.** Effect of abnormality type [**(A)** and **(D)**], translocation category **(B)** and translocation breakpoint position **(C)** are shown. Statistically significant levels are as follows: ***P<0.001, **P<0.01 and *P<0.05.

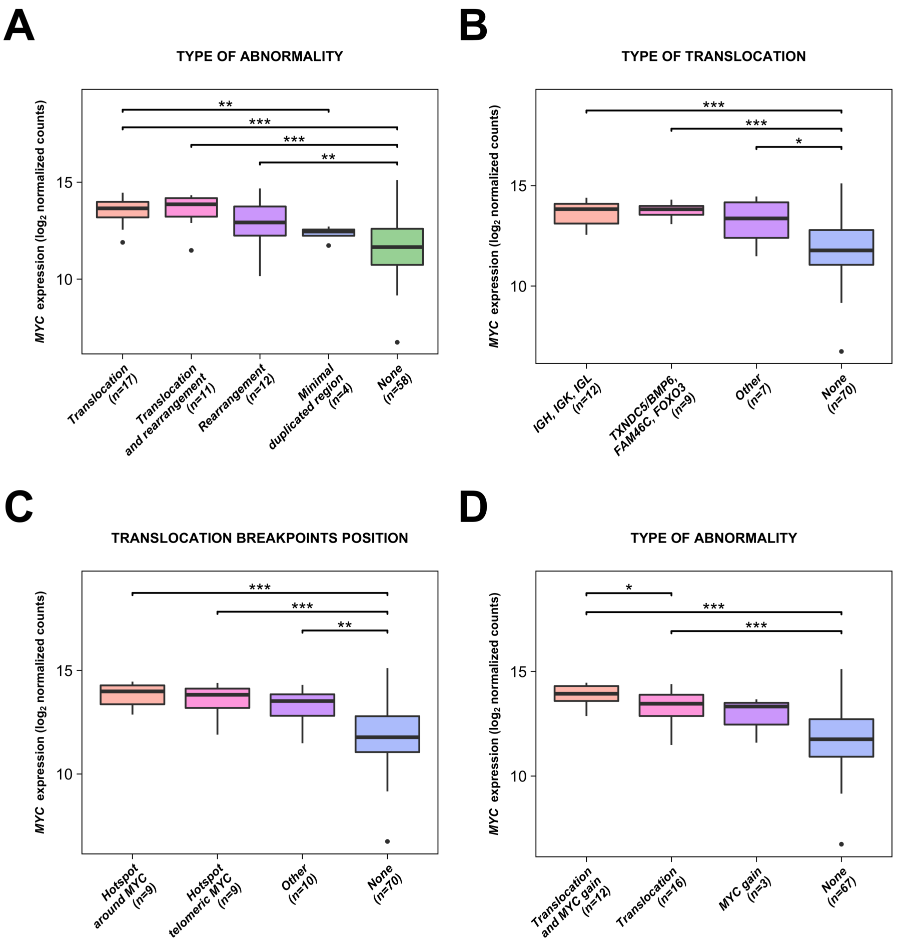

**Supplementary Figure 7: Copy-number abnormalities analysis at 8q24. (A)** Copy-number gains excluding tandem-duplications with two minimal gained regions. **(B)** Tandem-duplications with one minimal tandem-duplicated region. **(C)** Losses excluding deletions with one minimal lost region. **(D)** Deletions with two minimal deleted regions. Tandem-duplication and deletions were tested by paired-end read based analysis in a dataset of 1249 cases. Losses and gains were analyzed using tumor/control ratio depth analysis in a dataset of 97 cases with targeted sequencing. Total of three and 18 cases with complex intra-chromosomal rearrangement (more than five rearrangements) were excluded from analysis. Position of *MYC* (red) and other genes (gray) is shown.

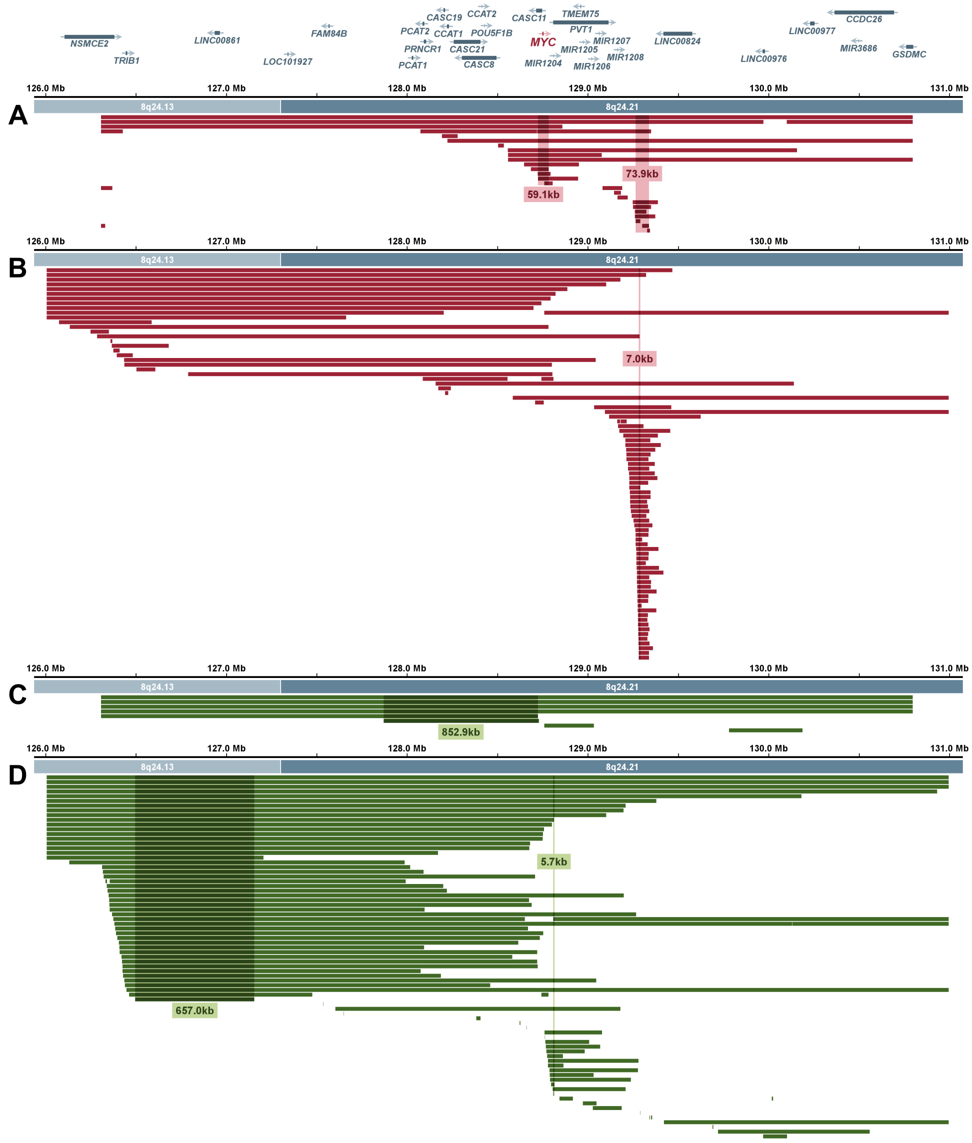

**Supplementary Figure 8: Frequency of copy-number abnormalities per position in *MYC* region.** Gains (red)/losses (green) are shown in upper part and tandem-duplications (red)/deletions (green) are shown in lower part. Tandem-duplication and deletions were tested by paired-end read based analysis in a dataset of 1249 cases. Losses and gains were analyzed using tumor/control ratio depth analysis in a dataset of 97 cases with targeted sequencing. Total of three and 18 cases with complex intra-chromosomal rearrangement (more than five rearrangements) were excluded from analysis. Position of *MYC* (red) and other genes (gray) is shown.

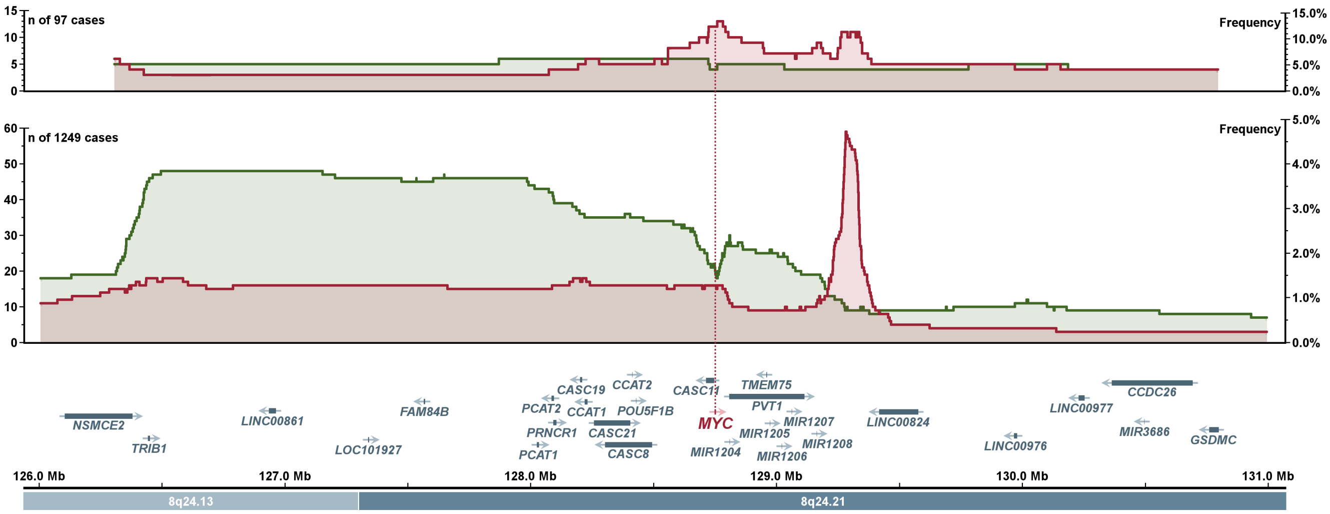

**Supplementary Figure 9: RNA-sequencing expression analysis of *MYC* and *PVT1* in relation to chromosomal abnormalities at 8q24 in hyperdiploidy group.** Effect of abnormality type [**(A)** and **(D)**], translocation category [**(B)** and **(E)**] and translocation breakpoint position [**(C)** and **(F)**] are shown for *MYC* and *PVT1*, respectively. Statistically significant levels are as follows: ***P<0.001, **P<0.01 and *P<0.05.

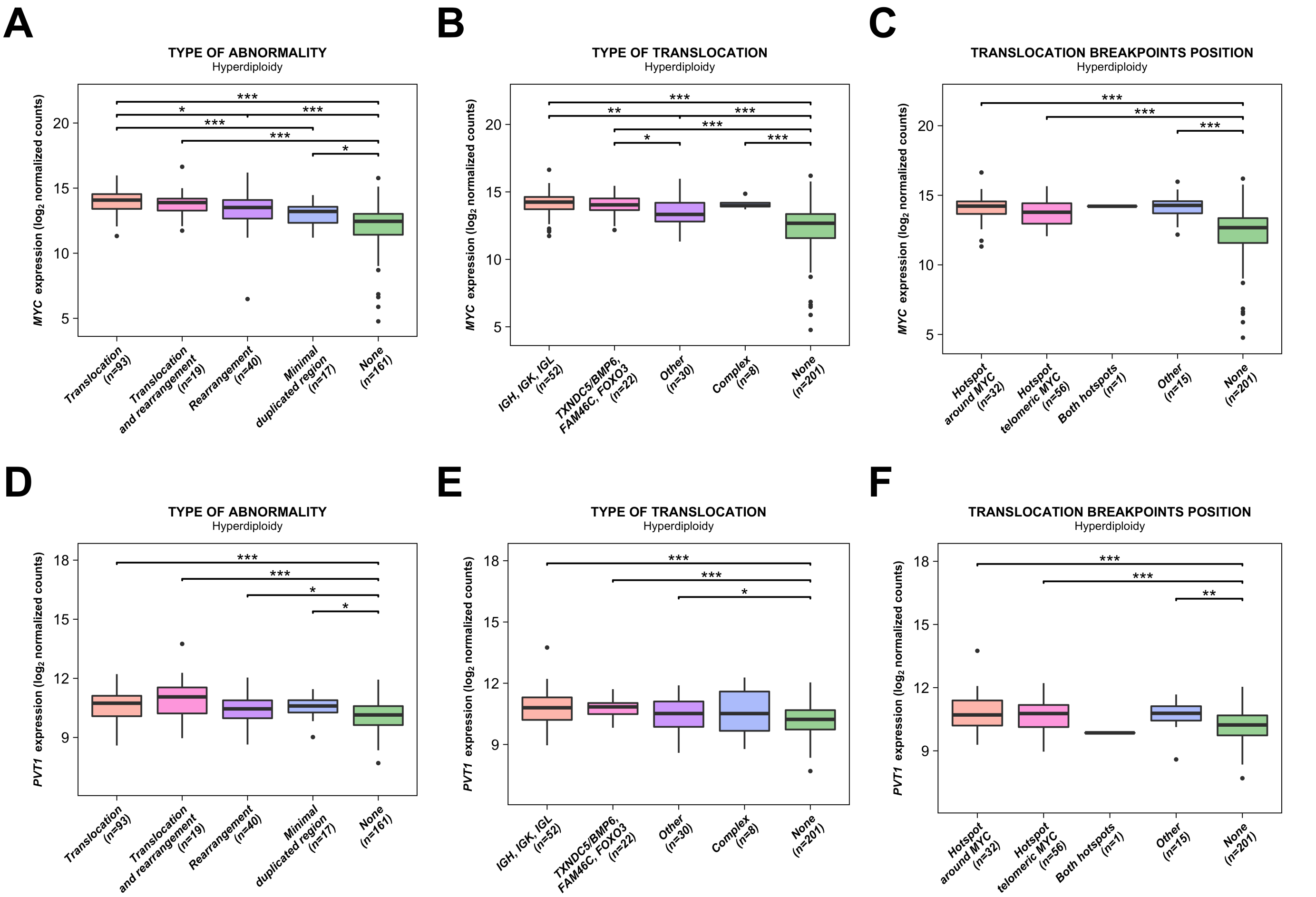

**Supplementary Figure 10: RNA-sequencing expression analysis of *MYC* and *PVT1* in relation to chromosomal abnormalities at 8q24 in non-hyperdiploidy group.** Effect of abnormality type [**(A)** and **(D)**], translocation category [**(B)** and **(E)**] and translocation breakpoint position [**(C)** and **(F)**] are shown for *MYC* and *PVT1*, respectively. Statistically significant levels are as follows: ***P<0.001, **P<0.01 and *P<0.05.

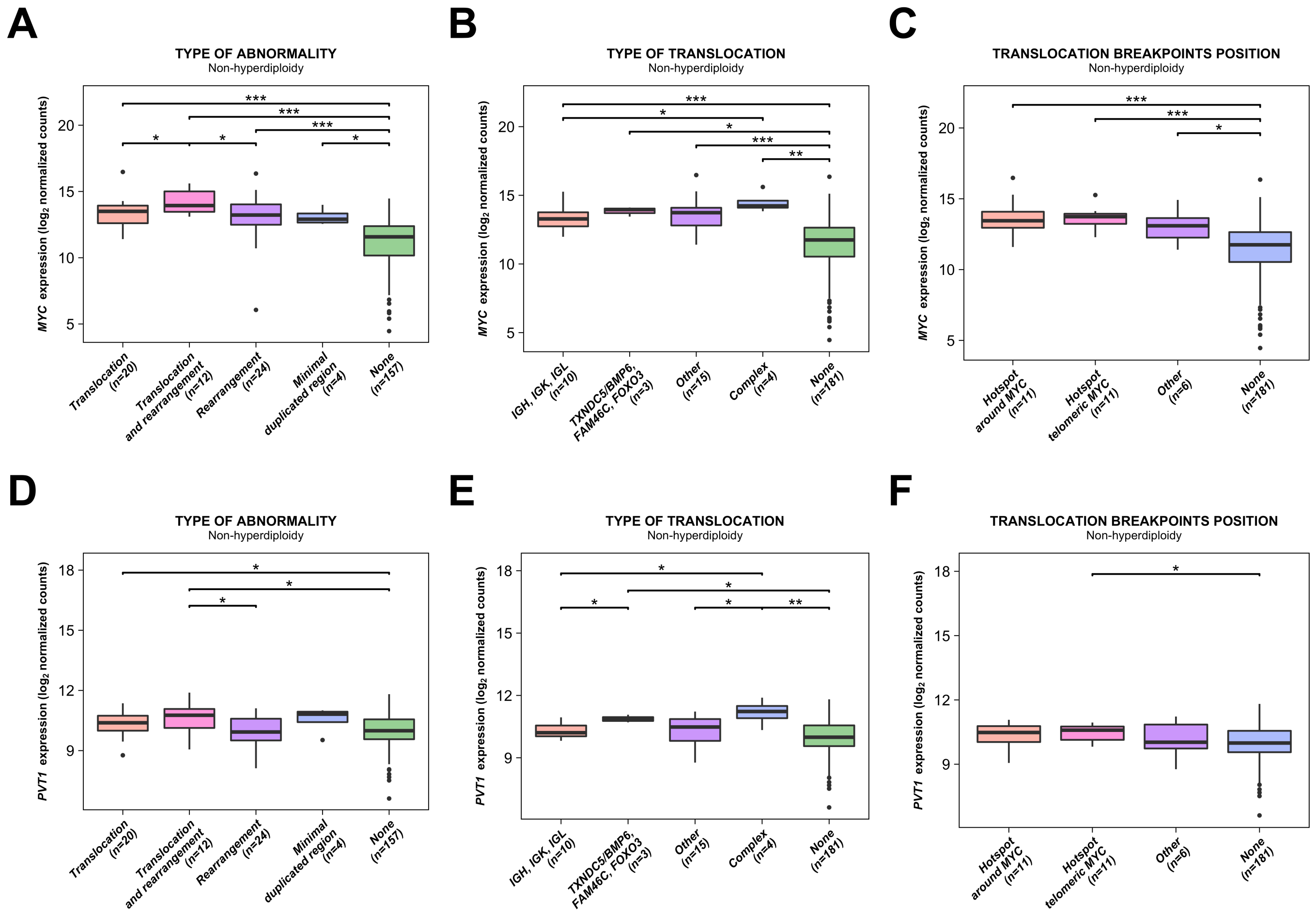

**Supplementary Figure 11: RNA-sequencing expression analysis of *MYC* and *PVT1* in relation to chromosomal abnormalities at 8q24 – comparison between hyperdiploidy and non-hyperdiploidy group.** Effect of abnormality type [**(A)** and **(D)**], translocation category [**(B)** and **(E)**] and translocation breakpoint position [**(C)** and **(F)**] are shown for *MYC* and *PVT1*, respectively. Statistically significant levels are as follows: ***P<0.001, **P<0.01 and *P<0.05.

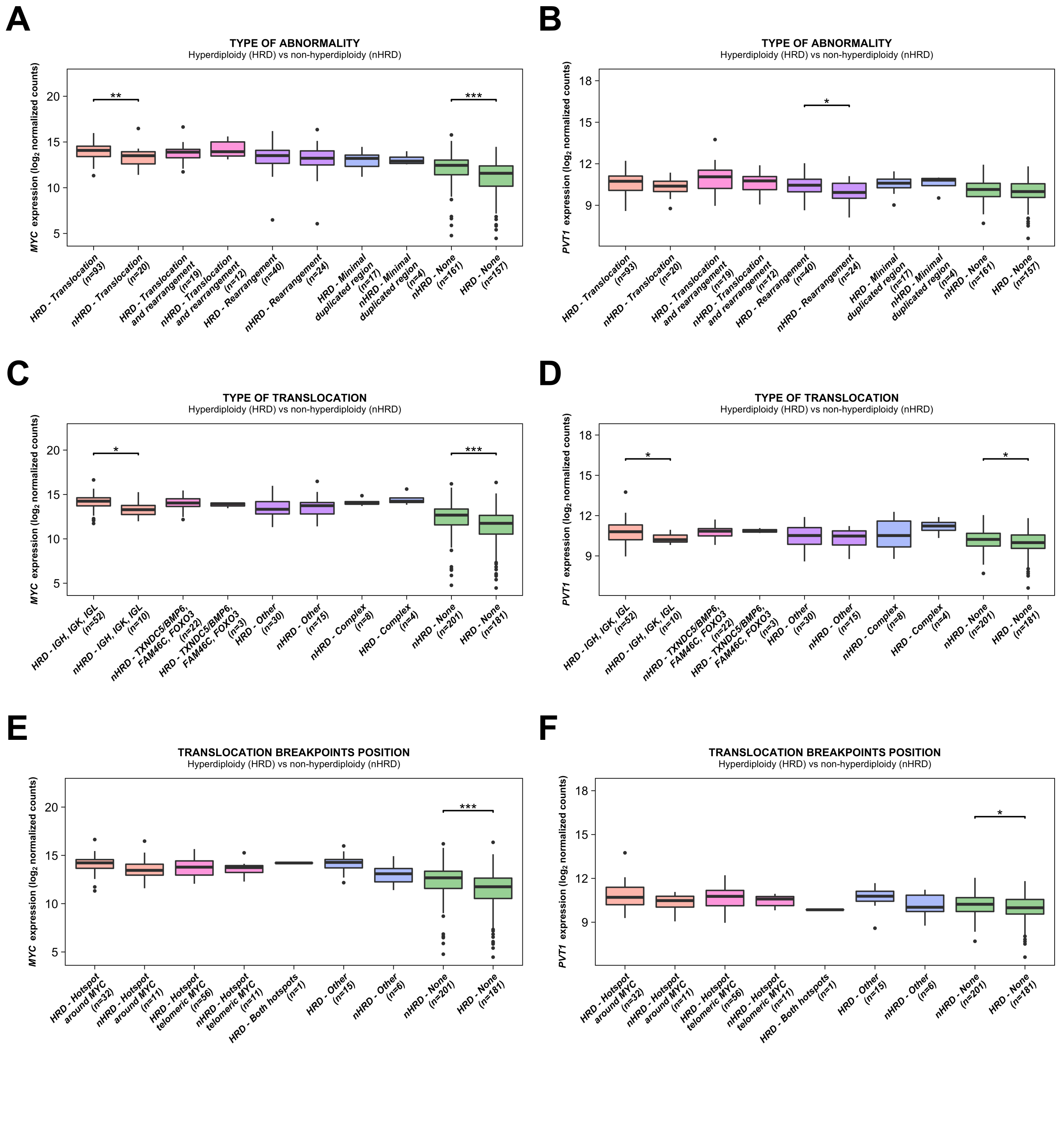

### Supplementary Alignments

**26425_RNAS_D-PL3539_CD138_KP-329MT**
**t(3;8)**

```
chr8:       GGGCACTTCTTGCTTTCTGCCTCCCATCAGTCATCCCAGGGGACGCCAGCTGCCACTTTG
            ||||||||||||||||||||||||||||| ||   |       |   |      ||
KP329:      GGGCACTTCTTGCTTTCTGCCTCCCATCATTCTCTCTGTCTCTCAGAATACTTAACACAT
               | | |     |  |    ||   |  |||||||||||||||||||||||||||||||
chr3:       TTCCTCCTGGAAGTACCCATATCTTTTACTTCTCTCTGTCTCTCAGAATACTTAACACAT
```

**37606_RNAS_42485_1-AS-RB-CD138-DNA_CD138_KP-084MT**
**t(2;8)**

```
chr8:       CTGCCAGAGATCCCTGTGTT-AACTGTGAACAGAGCCCTTTTCATCCTCTTGCATCAGAATTCCTG
            |||||||||||||||||||| ||  |    | |       ||||      |||       |  |||
KP084:      CTGCCAGAGATCCCTGTGTT-AATAGACTCCTGCCTGAACTTCAAGGCTATGCCCTGATGTCGCTG
            ||   | |||  |  ||||| |||||||||||||||||||||||||||||||||||||||||||||
chr2:       CTTAGAAAGAATCAAGTGTTGAATAGACTCCTGCCTGAACTTCAAGGCTATGCCCTGATGTCGCTG
```

**38738_RNAS_51065_1-AS-RB-CD138-DNA_CD138_KP-088MT**
**t(2;8)**

```
chr2:       CTGCTAGAGAGAGTTATGATCTCGCCACTGCACTCCACCCTGTGTGACAGAGTGAGACTC
            ||||||||||||||||||||           | | ||    || |        || | |
KP088:      CTGCTAGAGAGAGTTATGATGAGTGGGACCAAGTGCAATAGGTCTATGTCCAGGATAATT
            |            |   |||||||||||||||||||||||||||||||||||||||||||
chr8:       CATCCTTGACTCATCCAGATGAGTGGGACCAAGTGCAATAGGTCTATGTCCAGGATAATT
```

**24852_RNAS_D-PL3391_CD138_KP-214MT**
**t(3;8)**

```
chr3:       AAGTCTGATGTGATACCTACAAATTCAGCATATAAAATGAATCTAAAGAGTGCTTTGCT
            ||||||||||||||||||||||||||||||||||||    |   |  | |     |
KP214:      AAGTCTGATGTGATACCTACAAATTCAGCATATAAAGGACAGGCATTGGGGTTGCTTTG
              ||  |      |  |     ||  || ||||||||||||||||||||||||||||||
chr8:       TCGTTCGTAAACTTCACAGTTTATGAAGTATATAAAGGACAGGCATTGGGGTTGCTTTG
```

**27791_RNAS_D-PL3662_CD138_KP-141MT**
**t(8;19)**

```
chr19:      AATCACAGGCACATGCCATCATGCCTGGCTCTTTTTTTT
            ||||||||||||| |     |   |     |||
KP141:      AATCACAGGCACAGGAATGAAATTCATTTACTTAAAAAG
            ||  |     |||||||||||||||||||||||||||||
chr8:       AAGAATGACTACAGGAATGAAATTCATTTACTTAAAAAG
```

**35250_RNAS_D-PL4968_21925_1-AS-RB-CD138-DNA_CD138_KP-232MT**
**t(6;8)**

```
Chr8:       AGATTATAACCTTTTTAGGAGCAGCACACATTGTACTTACTATATCTTTTTATC
            ||||||||||||||||||||||||||||||   |     ||        ||||
KP232:      AGATTATAACCTTTTTAGGAGCAGCACACAGGTTCTCACCTTCTGTGGCTTATT
            |    | |||                ||||||||||||||||||||||||||||
Chr6:       AAGAGACAACTGACAACCCGAACTTCCACAGGTTCTCACCTTCTGTGGCTTATT
```

**39017_RNAS_55603_1-AS-RB-CD138-DNA_CD138_KP-305MT**
**t(6;8)**

```
Chr8:       GTGGGTCTTCATGATGTTCTTGTGATAGTGAGTAAGTCTC
            |||||||||||| |    ||     |      |
KP305:      GTGGGTCTTCATCAGCTCCTCTCTTTCTCTCCTGCACAAG
            |    | ||| |||||||||||||||||||||||||||||
Chr6:       GGCTTTGTTCTTCAGCTCCTCTCTTTCTCTCCTGCACAAG
```

**35830_RNAS_29095_1-AS-RB-CD138-DNA_CD138_KP-157MT**
**t(6;8)**

```
Chr8:       GACTAATACTCTTTTACCTAATCAGAGCCTGGCATGGTGCAGGTATATGAAATGA
            |||||||||||||||||||||||||||||||||  | ||    |
KP157:      GACTAATACTCTTTTACCTAATCAGAGCCTGGCCGGCTGATTCTCGGGTTGTGCC
              |  |  |     |          ||    ||||||||||||||||||||||||
Chr6:       CCCCCAGCCCCAGCTCTGGCCCTGCAGAAATGCCGGCTGATTCTCGGGTTGTGCC
```

### Supplementary Tables

#### Supplementary Table 1: Patients datasets characteristics, techniques for analysis and available number of samples.

| | Overall (n=1280) | UAMS (n=100) | UK (n=461) | MMRF (n=706) |
|---|---|---|---|---|
| | **Data availability and method** | | | |
| **Structural changes*** | n=1267/1267 (100%) | TS<br>n=100/100 (100%) | WES***<br>n=461/461 (100%) | WGS<br>n=706/706 (100%) |
| **NS-SNVs** | n=1264/1267 (99.8%) | TS<br>n=100/100 (100%) | WES***<br>n=461/461 (100%) | WES<br>n=703/706 (99.6%) |
| **CNAs**** | n=100/1267 (7.9%) | TS<br>n=100/100 (100%) | NA | NA |
| **Gene expression** | n=669/1267 (52.8%) | Microarray<br>n=98/100(98.0%) | NA | RNA-Seq<br>n=571/706 (80.9%) |
| **Metadata** | n=1262/1267 (99.6%) | n=98/100 (98.0%) | n=461/461 (100%) | n=703/706 (99.6%) |
| | **Basic characteristics** | | | |
| **Median Age [years] (range)** | 65.0 (30.4-93.0) | 60.7 (30.4-75.2) | 68.0 (31.0-89.0) | 64.0 (31.0-93.0) |
| **Age >= 65 years** | 672/1262 (53.2%) | 27/98 (27.6%) | 299/461 (64.9%) | 346/703 (49.2%) |
| **ISS stage 1** | 370/1214 (30.5%) | 28/98 (28.6%) | 105/436 (24.1%) | 237/680 (34.9%) |
| **ISS stage 2** | 456/1214 (37.6%) | 41/98 (41.8%) | 169/436 (38.8%) | 246/680 (36.2%) |
| **ISS stage 3** | 388/1214 (32.0%) | 29/98 (29.6%) | 162/436 (37.2%) | 197/680 (29.0%) |
| **t(4;14)** | 156/1262 (12.4%) | 10/98 (10.2%) | 58/461 (12.6%) | 88/703 (12.5%) |
| **t(6;14)** | 19/1262 (1.5%) | 6/98 (6.1%) | 5/461 (1.1%) | 8/703 (1.1%) |
| **t(8;14)** | 7/1164 (0.6%) | ND | 1/461 (0.2%) | 6/703 (0.9%) |
| **t(11;14)** | 237/1262 (18.8%) | 12/98 (12.2%) | 87/461 (18.9%) | 138/703 (19.6%) |
| **t(14;16) or t(14;20)** | 65/1262 (5.2%) | 6/98 (6.1%) | 20/461 (4.3%) | 39/703 (5.5%) |

*Translocations and chromosomal rearrangements including deletions, inversions and tandem-duplications

**Copy-number abnormalities analyzed by tumor/control depth ratio

**With custom enrichment for *MYC* region

**Supplementary Table 2: List of *MYC* non-synonymous variant in a dataset of 1264 myeloma patients.**

| n | Protein level | cDNA level | Type of variant | PROVEAN/SIFT prediction |
|---|---|---|---|---|
| 1 | p.Ser6Arg | c.18C>G | Missense mutation | Neutral/Damaging |
| 1 | p.Pro43fs | c.124delC | Frame-shift deletion | NA/NA |
| 1 | p.Ala44Val | c.131C>T | Missense mutation | Neutral/Damaging |
| 1 | p.Pro60Ser | c.178C>T | Missense mutation | Deleterious/Damaging |
| 1 | p.Val77fs | c.229dupG | Frame-shift insertion | NA/NA |
| 2 | p.Ser146Leu | c.437C>T | Missense mutation | Deleterious/Damaging |
| 1 | p.Val280del | c.834_836delTGT | In-frame deletion | Deleterious/NA |
| 1 | p.Ser420Tyr | c.1259C>A | Missense mutation | Deleterious/Damaging |

**Supplementary Table 3: Frequency of *MYC* translocation in datasets of 100 patients with targeted sequencing (TS), 461 patients with whole exome sequencing (WES) and 706 patients with whole genome sequencing (WGS).**

| Dataset | Translocation | Intra-locus rearrangement | Translocation and/or intra-locus rearrangement |
|---|---|---|---|
| TS | 29.0% (29/100) | 23.0% (23/100) | 41.0% (41/100) |
| WES | 23.6% (109/461) | 12.8% (59/461) | 32.8% (151/461) |
| WGS | 25.6% (181/706) | 16.4% (116/706) | 37.4% (264/706) |
| **COMBINED** | **25.2% (319/1267)** | **15.6% (198/1267)** | **36.0% (456/1267)** |

**Supplementary Table 4: Proportion of number of chromosomes involved in *MYC* translocation in datasets of 100 patients with targeted sequencing (TS), 461 patients with whole exome sequencing (WES) and 706 patients with whole genome sequencing (WGS).**

| Dataset | n=2 | n=3 | n=4 | n=5 | n>5 |
|---|---|---|---|---|---|
| TS | 69.0% (20/29) | 24.1% (7/29) | 6.9% (2/29) | 0.0% (0/29) | 0.0% (0/29) |
| WES | 76.1% (83/109) | 18.3% (20/109) | 5.5% (6/109) | 0.0% (0/109) | 0.0% (0/109) |
| WGS | 52.5% (95/181) | 25.4% (46/181) | 9.9% (18/181) | 4.4% (8/181) | 7.7% (14/181) |
| **COMBINED** | **62.1% (198/319)** | **22.9% (73/319)** | **8.2% (26/319)** | **2.5% (8/319)** | **4.4% (14/319)** |

**Supplementary Table 5: List of *MYC* translocation partners present in at least five cases in the dataset of 1253 non-complex NDMM patients.**

| Chromosome band | Position | Size, Mb | Frequency | Super-enhancer--associated genes in MM.1S cell line[1] | Immunoglobulin gene locus | Overlapped high expressed genes* | Candidate genes involved in *MYC* deregulation |
|---|---|---|---|---|---|---|---|
| **14q32.33** | chr14:105013903-107220085 | 2.2 | 5.0% (63/1253) | *MYC*[†], *TMEM121* | *IGH* | *SIVA1, AKT1, MTA1, IGHG2, IGHA1, IGHG1* | ***IGH*** |
| **22q11.22/22q11.23** | chr22:22658283-24193029 | 1.5 | 5.0% (63/1253) | *IGLL5, DERL3, LOC284889, MIF, MIR650, SLC2A11* | *IGL* | *IGLL5, IGLC1, IGLC2, BCR, SMARCB1, DERL3* | ***IGL*** |
| **6p24.3** | chr6:7727323-8387494 | 0.7 | 2.7% (34/1253) | *BMP6, MUTED-TXNDC5, TXNDC5, EEF1E1-MUTED, PIP5K1P1* | - | *BMP6, TXNDC5* | ***BMP6 TXNDC5*** |
| **2p11.2** | chr2:88858600-90253854 | 1.4 | 2.1% (26/1253) | - | *IGK* | *EIF2AK3, ANKRD36BP2, IGKC* | ***IGK*** |
| **1p12** | chr1:118158927-118431479 | 0.3 | 1.6% (20/1253) | *FAM46C* | - | *FAM46C* | ***FAM46C*** |
| **6q21** | chr6:108876006-109352787 | 0.5 | 1.1% (14/1253) | *FOXO3* | - | *FOXO3* | ***FOXO3*** |
| **11q13.4** | chr11:72732494-73092358 | 0.4 | 0.7% (9/1253) | - | - | *FCHSD2* | ***FCHSD2*** |
| **11q13.3** | chr11:68923361-69978263 | 1.1 | 0.6% (8/1253) | - | *IGH* associated | *CCND1*[‡] | ***IGH***[§] |
| **2p14** | chr2:64365459-66730504 | 2.4 | 0.5% (6/1253) | *SERTAD2, LOC339807* | - | *PELI1, AFTPH, SERTAD2, SLC1A4, RAB1A, ACTR2* | ***SERTAD2*** |
| **8q23.3** | chr8:113454929-115844684 | 2.4 | 0.5% (6/1253) | - | - | - | **unknown** |
| **4q31.3** | chr4:153354954-153619440 | 0.3 | 0.4% (5/1253) | - | - | *FBXW7* | ***FBXW7*** |
| **13q22.3** | chr13:78500741-78766726 | 0.3 | 0.4% (5/1253) | - | - | *MYCBP2* (in <1Mb distance) | ***MYCBP2*** |

* >95% of 571 patients tested by RNA-seq show $\log_2$ normalized counts >10; † Due to the translocation t(8;14) in MM.1S; ‡ In subgroup of patients with t(11;14); § All 8 patients show t(11;14); ‖ Loven *et al.* 2013.

References:

1. Loven J, Hoke HA, Lin CY, *et al.* Selective inhibition of tumor oncogenes by disruption of super-enhancers. Cell. 2013;153(2):320-334.

**Supplementary Table 6: List of *MYC* translocation partners.** n = number of cases in the dataset of 1253 non-complex patients.

| Chromosomal band | Genome position | Size, bp | n |
|---|---|---|---|
| 1p35.3/1p36.11 | chr1:27739863-28392188 | 652325 | 2 |
| 1p34.3 | chr1:35894394-35894431 | 37 | 1 |
| 1p34.2 | chr1:40488361-40756958 | 268597 | 2 |
| 1p32.3 | chr1:52981834 | --- | 1 |
| 1p31.3 | chr1:66791014-66800178 | 9164 | 1 |
| 1p22.3 | chr1:85996952 | --- | 1 |
| 1p22.2 | chr1:88948039 | --- | 1 |
| 1p12 | chr1:118158927-118431479 | 272552 | 20 |
| 1q21.3 | chr1:150651915 | --- | 1 |
| 1q23.3 | chr1:161726404-163255527 | 1529123 | 3 |
| 1q25.2 | chr1:178531670-178583707 | 52037 | 1 |
| 1q25.3 | chr1:184720646 | --- | 1 |
| 1q32.1 | chr1:203051534-203274522 | 222988 | 2 |
| 2p23.3 | chr2:25549035-27405899 | 1856864 | 2 |
| 2p16.2 | chr2:54318733-54778431 | 459698 | 2 |
| 2p14 | chr2:64365459-66730504 | 2365045 | 6 |
| 2p13.3 | chr2:70401712 | --- | 1 |
| 2p11.2 | chr2:88858600-90253854 | 1395254 | 26 |
| 2q21.2 | chr2:134989575 | --- | 1 |
| 2q24.3 | chr2:166415936 | --- | 1 |
| 2q31.1 | chr2:173424877-173425255 | 378 | 1 |
| 2q32.1/2q32.2 | chr2:188778708-189976753 | 1198045 | 2 |
| 3p22.3 | chr3:32207945-32208479 | 534 | 1 |
| 3p21.31 | chr3:46330235-46395429 | 65194 | 1 |
| 3p21.31 | chr3:50196574-50389775 | 193201 | 1 |
| 3p21.1 | chr3:53079658 | --- | 1 |
| 3q13.2 | chr3:112244683-112249926 | 5243 | 1 |
| 3q26.2 | chr3:169228942-169495458 | 266516 | 1 |
| 3q26.31 | chr3:171762090 | --- | 1 |
| 4p16.3 | chr4:1857489-2732041 | 874552 | 2 |
| 4p15.2 | chr4:25908245-25908304 | 59 | 1 |
| 4q31.21 | chr4:141697852 | --- | 1 |
| 4q31.3 | chr4:153354954-153619440 | 264486 | 5 |
| 4q34.3 | chr4:179807156 | --- | 1 |
| 4q35.1 | chr4:185454754-185622688 | 167934 | 1 |

| | | | |
|---|---|---|---|
| 5p14.3/5p15.1 | chr5:17929865-19465553 | 1535688 | 1 |
| 5q11.2 | chr5:55401472 | --- | 1 |
| 5q14.3 | chr5:88450919-88796074 | 345155 | 3 |
| 5q22.1 | chr5:109858587-109859680 | 1093 | 1 |
| 5q31.2 | chr5:139433830 | --- | 1 |
| 5q33.1 | chr5:149829862-151044833 | 1214971 | 2 |
| 5q33.3 | chr5:156273797-156421100 | 147303 | 4 |
| 5q34 | chr5:160122942-160200603 | 77661 | 1 |
| 5q35.2 | chr5:173129032-173289854 | 160822 | 1 |
| 6p25.3 | chr6:194842-391218 | 196376 | 3 |
| 6p24.3 | chr6:7727323-8387494 | 660171 | 34 |
| 6p22.3 | chr6:21786639-23029400 | 1242761 | 1 |
| 6p21.2 | chr6:37035844-37546594 | 510750 | 2 |
| 6p21.1 | chr6:41858886-41993130 | 134244 | 1 |
| 6p12.1 | chr6:53795877-53961228 | 165351 | 1 |
| 6q15 | chr6:88632133-89925597 | 1293464 | 2 |
| 6q21 | chr6:106041941-107122607 | 1080666 | 4 |
| 6q21 | chr6:108876006-109352787 | 476781 | 14 |
| 6q22.31 | chr6:119693342 | --- | 1 |
| 7p21.3 | chr7:7914145-7999977 | 85832 | 1 |
| 7p21.3 | chr7:11159783-11593112 | 433329 | 1 |
| 7p21.2 | chr7:13962197-13967016 | 4819 | 1 |
| 7p15.2 | chr7:26008444-26149402 | 140958 | 2 |
| 7p14.3 | chr7:34559624 | --- | 1 |
| 7q21.12 | chr7:87048847-87049154 | 307 | 1 |
| 7q22.3 | chr7:105450486-105475544 | 25058 | 1 |
| 7q31.33/7q32.1 | chr7:126925518-127156534 | 231016 | 1 |
| 7q33 | chr7:137678731-137683581 | 4850 | 1 |
| 7q34 | chr7:139455673-139639290 | 183617 | 2 |
| 8p12 | chr8:29407990 | --- | 1 |
| 8q21.11 | chr8:77422720 | --- | 1 |
| 8q21.13 | chr8:83875044 | --- | 1 |
| 8q21.3 | chr8:87616372 | --- | 1 |
| 8q21.3 | chr8:90568607 | --- | 1 |
| 8q22.1 | chr8:95826016 | --- | 1 |
| 8q22.1 | chr8:98499039-98652529 | 153490 | 3 |
| 8q22.2 | chr8:101487464 | --- | 1 |
| 8q22.3 | chr8:103285796-105762724 | 2476928 | 3 |
| 8q23.1 | chr8:106251133 | --- | 1 |

| | | | |
|---|---|---|---|
| 8q23.2 | chr8:111993713 | --- | 1 |
| 8q23.3 | chr8:113454929-115844684 | 2389755 | 6 |
| 8q24.12 | chr8:119641772-120883792 | 1242020 | 2 |
| 8q24.22 | chr8:132435708-132587826 | 152118 | 1 |
| 8q24.3 | chr8:141922737 | --- | 1 |
| 8q24.3 | chr8:145287815 | --- | 1 |
| 9q21.13 | chr9:79164605-79190341 | 25736 | 1 |
| 9q22.2 | chr9:93446606-93817496 | 370890 | 1 |
| 9q34.11/9q34.13 | chr9:132193351-134167408 | 1974057 | 2 |
| 10p14 | chr10:6742071-6868227 | 126156 | 1 |
| 10q22.3 | chr10:79061510-79615885 | 554375 | 1 |
| 10q24.32 | chr10:104146091-104159683 | 13592 | 1 |
| 10q25.2 | chr10:113570048 | --- | 1 |
| 10q26.13 | chr10:125148066-125191492 | 43426 | 1 |
| 11p15.1 | chr11:19131381-19132109 | 728 | 1 |
| 11p12 | chr11:36704013-37514028 | 810015 | 1 |
| 11p11.2 | chr11:44998507-45076079 | 77572 | 1 |
| 11q12.1 | chr11:58882384-58896224 | 13840 | 1 |
| 11q13.2 | chr11:66775196-66958643 | 183447 | 1 |
| 11q13.3 | chr11:68923361-69978263 | 1054902 | 8 |
| 11q13.4 | chr11:72732494-73092358 | 359864 | 9 |
| 11q14.1 | chr11:82400131-82859140 | 459009 | 1 |
| 11q22.1 | chr11:98859287 | --- | 1 |
| 11q23.3 | chr11:118938634-119240262 | 301628 | 1 |
| 11q24.3 | chr11:128243023-128717432 | 474409 | 2 |
| 11q25 | chr11:131995275 | --- | 1 |
| 12p13.32 | chr12:3835029-4707543 | 872514 | 3 |
| 12p11.23 | chr12:26941957 | --- | 1 |
| 12p11.21 | chr12:32065050 | --- | 1 |
| 12q13.11 | chr12:47758587 | --- | 1 |
| 12q14.1 | chr12:58147672-58175116 | 27444 | 1 |
| 12q15 | chr12:68868290-68889029 | 20739 | 1 |
| 12q22 | chr12:93122961 | --- | 1 |
| 12q23.3 | chr12:105207790-105282218 | 74428 | 1 |
| 13q13.3 | chr13:35705874-35706246 | 372 | 1 |
| 13q14.2 | chr13:48924610 | --- | 1 |
| 13q22.3 | chr13:78500741-78766726 | 265985 | 5 |
| 14q23.3 | chr14:65720336-65901445 | 181109 | 2 |
| 14q24.2 | chr14:72818022 | --- | 1 |

| | | | |
|---|---|---|---|
| 14q32.12 | chr14:92883617 | --- | 1 |
| 14q32.33 | chr14:105013903-107220085 | 2206182 | 63 |
| 15q11.2 | chr15:24141512 | --- | 1 |
| 15q13.3 | chr15:31676325 | --- | 1 |
| 15q22.31 | chr15:64018516 | --- | 1 |
| 15q24.1/15q24.2 | chr15:75096204-75459662 | 363458 | 2 |
| 15q25.1 | chr15:80357809-81585731 | 1227922 | 2 |
| 16p11.2 | chr16:29229191-33437673 | 4208482 | 4 |
| 16q23.1/16q23.2 | chr16:78569426-79239195 | 669769 | 4 |
| 17p13.2 | chr17:3624808-4455123 | 830315 | 2 |
| 17p13.1 | chr17:8214064 | --- | 1 |
| 17p11.2 | chr17:16730207-20401949 | 3671742 | 1 |
| 17q12 | chr17:32532251-32536335 | 4084 | 1 |
| 17q21.32 | chr17:45205683-45354778 | 149095 | 1 |
| 17q23.3 | chr17:62391611-62491820 | 100209 | 2 |
| 17q25.2 | chr17:75110491-75118485 | 7994 | 1 |
| 18q21.33 | chr18:60774868-60841160 | 66292 | 1 |
| 19p13.3 | chr19:1559331-2555073 | 995742 | 4 |
| 19p13.2 | chr19:12804957-13638653 | 833696 | 1 |
| 19p13.11/19p13.12 | chr19:16247562-16599708 | 352146 | 2 |
| 19q13.32 | chr19:46136027 | --- | 1 |
| 19q13.33 | chr19:48289453-49745349 | 1455896 | 1 |
| 20p12.2 | chr20:9882367-10263026 | 380659 | 1 |
| 20p12.1 | chr20:17819354 | --- | 1 |
| 20q11.21 | chr20:30814222-30911114 | 96892 | 1 |
| 20q11.22 | chr20:32435299-32623087 | 187788 | 3 |
| 20q11.22 | chr20:34171969-34271613 | 99644 | 1 |
| 20q13.12/20q13.13 | chr20:45975160-47499328 | 1524168 | 4 |
| 20q13.13 | chr20:49076881-49165445 | 88564 | 1 |
| 21q11.2/21q21.1 | chr21:15408264-16910043 | 1501779 | 1 |
| 21q22.3 | chr21:44751330-44832028 | 80698 | 1 |
| 22q11.22/22q11.23 | chr22:22658283-24193029 | 1534746 | 63 |
| 22q12.1 | chr22:28457893-29210349 | 752456 | 2 |
| 22q13.1/22q13.2 | chr22:40604732-42203185 | 1598453 | 3 |
| Xq28 | chrX:153123577-153313715 | 190138 | 1 |
| Un_gl000220unk | chrUn_gl000220:115792-145495 | 29703 | 3 |

### Supplementary Table 7: Genes deregulated with *MYC* abnormalities.

| Gene symbol | Location | Full name | Regulation* | GEN foldchg | GEN FDR | EPX foldchg | EPX FDR | MYC motif | Evidence |
|---|---|---|---|---|---|---|---|---|---|
| **UP-REGULATED** | | | | | | | | | |
| ***MYC*** | 8q24.21 | *MYC* proto-oncogene, bHLH transcription factor | GEN/EXP | 3.5 | 7.1E-28 | 5.9 | 1.6E-67 | | Up-regulation confirmed in most of the studies |
| ***HK2*** | 2p12 | hexokinase 2 | GEN/EXP | 3.6 | 1.7E-18 | 2.4 | 2.3E-08 | | Validated *MYC* target genes.[3]<br>Up-regulated genes selected in supervised analyses to discriminate cells expressing *MYC* from control cells expressing GFP.[4]<br>Genes up-regulated in P493-6 cells (B lymphocyte, Burkitt's lymphoma) by *MYC* and down-regulated by the combination of *MYC* and serum.[5] |
| ***LAMP5*** | 20p12.2 | lysosomal associated membrane protein family member 5 | GEN/EXP | 3.4 | 4.0E-07 | 2.2 | 2.4E-03 | | |
| ***CGREF1*** | 2p23.3 | cell growth regulator with EF-hand domain 1 | GEN/EXP | 2.9 | 1.4E-10 | 2.7 | 3.6E-09 | YES[1,2] | Genes up-regulated in primary epithelial breast cancer cell culture over-expressing *MYC* gene.[4] |
| ***DDN*** | 12q13.12 | dendrin | GEN/EXP | 2.5 | 1.6E-26 | 2.5 | 5.4E-26 | | |
| ***SNHG4*** | 5q31.2 | small nucleolar RNA host gene 4 | GEN/EXP | 2.5 | 5.1E-24 | 2.4 | 2.7E-22 | | |
| ***SORD*** | 15q21.1 | sorbitol dehydrogenase | GEN/EXP | 2.3 | 1.6E-26 | 2.2 | 2.2E-25 | YES[2] | Validated *MYC* target genes.[3]<br>Up-regulated genes selected in supervised analyses to discriminate cells expressing *MYC* from control cells expressing GFP.[4]<br>Genes up-regulated in primary epithelial breast cancer cell culture over-expressing *MYC* gene.[4]<br>Genes up-regulated in P493-6 cells (B lymphocyte, Burkitt's lymphoma) induced to express *MYC*.[6]<br>Genes up-regulated in K562 cells (lymphoblast, chronic myelogenous leukemia) by *MYC* in the presence of *CKN1B*.[7]<br>Genes directly up-regulated in P493-6 cells (B lymphocyte, Burkitt's lymphoma) by *MYC*.[8] |
| ***STEAP3*** | 2q14.2 | STEAP3 metalloreductase | GEN/EXP | 2.1 | 1.3E-10 | 2.3 | 2.9E-13 | | |
| ***SPTBN2*** | 11q13.2 | spectrin beta, non-erythrocytic 2 | GEN/EXP | 2.3 | 2.7E-14 | 2.1 | 8.4E-12 | | |
| ***VPS9D1-AS1*** | 16q24.3 | VPS9D1 antisense RNA 1 | GEN/EXP | 2.0 | 5.1E-24 | 2.2 | 4.0E-29 | | |
| ***HPDL*** | 1p34.1 | 4-hydroxyphenylpyruvate dioxygenase like | GEN/EXP | 2.1 | 1.4E-15 | 2.0 | 1.0E-13 | | |
| ***RPH3A*** | 12q24.13 | rabphilin 3A | GEN/EXP | 2.3 | 5.2E-07 | 1.8 | 1.3E-03 | | |
| ***ANKRD13B*** | 17q11.2 | ankyrin repeat domain 13B | GEN/EXP | 2.0 | 2.3E-22 | 2.0 | 2.9E-24 | YES[2] | |
| ***SLC19A1*** | 21q22.3 | solute carrier family 19 member 1 | GEN/EXP | 2.0 | 1.8E-23 | 1.9 | 8.3E-20 | | Validated *MYC* target genes.[3]<br>Up-regulated genes selected in supervised analyses to discriminate cells expressing *MYC* from control cells expressing GFP.[4]<br>Genes up-regulated in primary epithelial breast cancer cell culture over-expressing *MYC* gene.[4]<br>Targets of *MYC* identified by ChIP on chip in cultured cell lines, focusing on E-box-containing genes; high affinity bound subset.[9]<br>Genes identified by ChIP within the high-affinity group of *MYC* targets.[10]<br>Genes whose promoters are bound by *MYC*, according to *MYC* Target Gene Database.[11]<br>Genes directly up-regulated in P493-6 cells (B lymphocyte, Burkitt's lymphoma) by *MYC*.[8] |
| ***SSTR3*** | 22q13.1 | somatostatin receptor 3 | GEN/EXP | 2.1 | 2.8E-04 | 1.9 | 1.9E-03 | | |
| ***SEPT3*** | 22q13.2 | septin 3 | GEN/EXP | 2.2 | 1.9E-09 | 1.8 | 4.9E-05 | YES[1,2] | |
| ***EPHB4*** | 7q22.1 | EPH receptor B4 | GEN/EXP | 1.9 | 1.6E-23 | 1.9 | 1.3E-19 | | |
| ***MTHFD1L*** | 6q25.1 | methylenetetrahydrofolate dehydrogenase | GEN/EXP | 1.8 | 9.1E-11 | 2.0 | 6.4E-14 | | Genes directly up-regulated in P493-6 cells (B lymphocyte, Burkitt's lymphoma) by *MYC*.[8] |

| | | | | | | | | | |
|---|---|---|---|---|---|---|---|---|---|
| ***MFNG*** | 22q13.1 | MFNG O-fucosylpeptide 3-beta-N-acetylglucosaminyltransferase | GEN/EXP | 2.0 | 1.2E-08 | 1.8 | 3.0E-06 | | Targets of *MYC* and *MAX* identified by ChIP on chip in a Burkitt's lymphoma cell line; overlap set.[9] Genes whose promoters are bound by *MYC*, according to *MYC* Target Gene Database.[11] Genes directly up-regulated in P493-6 cells (B lymphocyte, Burkitt's lymphoma) by *MYC*.[8] |
| ***TMEM145*** | 19q13.2 | transmembrane protein 145 | GEN/EXP | 1.8 | 1.4E-13 | 2.0 | 1.2E-18 | | |
| ***HMCN2*** | 9q34.11 | hemicentin 2 | GEN/EXP | 2.0 | 3.1E-05 | 1.7 | 2.7E-03 | | |
| ***LRFN4*** | 11q13.2 | leucine rich repeat and fibronectin type III domain containing 4 | GEN/EXP | 1.8 | 2.6E-08 | 1.9 | 1.2E-10 | YES[2] | |
| ***GAS5*** | 1q25.1 | growth arrest specific 5 (non-protein coding) | GEN/EXP | 1.8 | 1.4E-18 | 1.9 | 1.6E-23 | | |
| ***CCDC78*** | 16p13.3 | coiled-coil domain containing 78 | GEN/EXP | 1.8 | 5.7E-13 | 1.8 | 2.2E-12 | | Up-regulated genes selected in supervised analyses to discriminate cells expressing *MYC* from control cells expressing GFP.[4] Genes up-regulated in primary epithelial breast cancer cell culture over-expressing *MYC* gene.[4] |
| ***SVOP*** | 12q24.11 | SV2 related protein | GEN/EXP | 2.1 | 2.0E-08 | 1.6 | 8.4E-04 | | |
| ***SCN3A*** | 2q24.3 | sodium voltage-gated channel alpha subunit 3 | GEN | 2.1 | 4.8E-06 | 1.4 | 7.7E-02 | | |
| ***ZC3HAV1L*** | 7q34 | zinc finger CCCH-type containing, antiviral 1 like | GEN/EXP | 1.6 | 1.7E-05 | 1.9 | 4.3E-09 | | |
| ***DIXDC1*** | 11q23.1 | DIX domain containing 1 | GEN/EXP | 1.9 | 1.2E-09 | 1.6 | 4.0E-05 | | |
| ***SLC43A1*** | 11q12.1 | solute carrier family 43 member 1 | GEN/EXP | 1.6 | 2.3E-05 | 1.8 | 2.6E-07 | YES[1,2] | Targets of *MYC* and *MAX* identified by ChIP on chip in a Burkitt's lymphoma cell line; overlap set.[9] Genes whose promoters are bound by *MYC*, according to *MYC* Target Gene Database.[11] |
| ***S1PR4*** | 19p13.3 | sphingosine-1-phosphate receptor 4 | GEN/EXP | 1.5 | 4.1E-02 | 1.9 | 7.2E-04 | | |
| ***ROR2*** | 9q22.31 | receptor tyrosine kinase like orphan receptor 2 | GEN/EXP | 1.9 | 1.1E-03 | 1.5 | 4.7E-02 | | Genes down-regulated after double Cre-lox knockout of both *APC* and *MYC* in small intestine.[12] Genes up-regulated after Cre-lox knockout of *APC* in the small intestine that require functional *MYC*.[12] Wnt target genes up-regulated after Cre-lox knockout of *APC* in the small intestine that require functional *MYC*.[12] |
| ***SEMA3G*** | 3p21.1 | semaphorin 3G | GEN/EXP | 1.4 | 5.6E-03 | 1.9 | 6.9E-08 | | |
| ***C4A*** | 6p21.33 | complement C4A (Rodgers blood group) | GEN | 2.1 | 1.0E-04 | 1.2 | 4.8E-01 | | |
| ***C4A-AS1*** | 6p21.33 | C4A antisense RNA 1 | GEN | 2.0 | 9.8E-07 | 1.2 | 2.8E-01 | | |
| ***C4B-AS1*** | 6p21.33 | C4B antisense RNA 1 | GEN | 2.0 | 9.8E-07 | 1.2 | 2.8E-01 | | |
| ***RELN*** | 7q22.1 | reelin | GEN | 2.0 | 3.5E-04 | 1.2 | 5.2E-01 | | |
| ***C4B*** | 6p21.33 | complement C4B | GEN | 2.0 | 3.0E-04 | 1.1 | 7.8E-01 | | |
| ***PTP4A3*** | 8q24.3 | protein tyrosine phosphatase type IVA, member 3 | GEN | 1.9 | 7.7E-03 | 1.1 | 7.1E-01 | | Genes down-regulated in hepatocellular carcinoma tissue of *MYC* and *TGFA* double transgenic mice.[13] |
| ***LDLRAD2*** | 1p36.12 | low density lipoprotein receptor class A domain containing 2 | GEN | 1.8 | 1.3E-04 | 1.2 | 5.1E-01 | | |
| **DOWN-REGULATED** | | | | | | | | | |
| ***MAGED4B*** | Xp11.22 | MAGE family member D4B | GEN/EXP | 2.6 | 1.5E-11 | 2.4 | 6.6E-09 | | |
| ***MAGED4*** | Xp11.22 | MAGE family member D4 | GEN/EXP | 2.6 | 7.6E-12 | 2.3 | 4.0E-09 | | |
| ***CD79A*** | 19q13.2 | CD79a molecule | GEN/EXP | 2.7 | 4.0E-08 | 1.9 | 7.0E-04 | | |
| ***CD28*** | 2q33.2 | CD28 molecule | GEN/EXP | 2.1 | 9.0E-06 | 2.5 | 9.4E-08 | | |
| ***PLEKHO1*** | 1q21.2 | pleckstrin homology domain containing O1 | GEN/EXP | 2.8 | 3.3E-14 | 1.7 | 4.3E-04 | | |
| ***NTNG1*** | 1p13.3 | netrin G1 | GEN/EXP | 2.5 | 7.6E-16 | 2.0 | 2.8E-08 | | |
| ***SCNN1B*** | 16p12.2 | sodium channel epithelial 1 beta subunit | GEN/EXP | 2.4 | 2.3E-14 | 2.1 | 6.5E-10 | | |
| ***CD27*** | 12p13.31 | CD27 molecule | GEN/EXP | 2.5 | 9.4E-09 | 2.0 | 3.2E-05 | | Genes down-regulated in P493-6 cells (B lymphocyte, Burkitt's lymphoma) by *MYC* and up-regulated by RNAi knockdown of *TFRC*.[14] |

| | | | | | | | | |
|---|---|---|---|---|---|---|---|---|
| ***MYADM*** | 19q13.42 | myeloid associated differentiation marker | GEN/EXP | 1.9 | 1.0E-03 | 2.5 | 5.4E-07 | |
| ***PTPRCAP*** | 11q13.2 | protein tyrosine phosphatase, receptor type C associated protein | GEN/EXP | 2.1 | 2.2E-04 | 2.3 | 6.7E-05 | |
| ***SLC22A17*** | 14q11.2 | solute carrier family 22 member 17 | GEN/EXP | 2.0 | 4.6E-04 | 2.4 | 3.7E-06 | |
| ***PPIC*** | 5q23.2 | peptidylprolyl isomerase C | GEN/EXP | 2.0 | 4.5E-06 | 2.3 | 1.0E-07 | |
| ***LAPTM5*** | 1p35.2 | lysosomal protein transmembrane 5 | GEN/EXP | 2.4 | 1.0E-05 | 1.8 | 8.4E-03 | Genes down-regulated in B cell lymphoma tumors expressing an activated form of *MYC*.[15]<br>Genes directly down-regulated in P493-6 cells (B lymphocyte, Burkitt's lymphoma) by *MYC*.[8] |
| ***LBH*** | 2p23.1 | limb bud and heart development | GEN/EXP | 2.2 | 8.4E-09 | 2.1 | 2.5E-07 | |
| ***RAP1GAP2*** | 17p13.3 | RAP1 GTPase activating protein 2 | GEN/EXP | 2.2 | 9.0E-13 | 2.0 | 4.7E-09 | |
| ***ARHGEF40*** | 14q11.2 | Rho guanine nucleotide exchange factor 40 | GEN/EXP | 1.9 | 1.1E-10 | 2.2 | 3.0E-15 | |
| ***TCN2*** | 22q12.2 | transcobalamin 2 | GEN/EXP | 2.0 | 7.3E-09 | 2.2 | 2.7E-11 | Genes down-regulated in K562 cells (lymphoblast, chronic myelogenous leukemia) expressing *TP53* and *MYC*.[16]<br>Genes down-regulated by *MYC*, according to the *MYC* Target Gene Database.[11] |
| ***BASP1*** | 5p15.1 | brain abundant membrane attached signal protein 1 | GEN/EXP | 1.9 | 3.3E-04 | 2.2 | 1.8E-05 | Genes directly down-regulated in P493-6 cells (B lymphocyte, Burkitt's lymphoma) by *MYC*.[8] |
| ***PCDHGC3*** | 5q31.3 | protocadherin gamma subfamily C, | GEN/EXP | 2.4 | 6.6E-10 | 1.6 | 4.3E-03 | |
| ***CXCL12*** | 10q11.21 | C-X-C motif chemokine ligand 12 | GEN/EXP | 2.0 | 2.7E-04 | 2.0 | 4.7E-04 | |
| ***MS4A1*** | 11q12.2 | membrane spanning 4-domains A1 | GEN/EXP | 2.0 | 3.5E-03 | 2.0 | 4.4E-03 | Genes down-regulated in B cell lymphoma tumors expressing an activated form of *MYC*.[15] |
| ***CNN3*** | 1p21.3 | calponin 3 | GEN/EXP | 2.2 | 3.4E-07 | 1.7 | 1.1E-03 | |
| ***SPRED1*** | 15q14 | sprouty related EVH1 domain containing 1 | GEN/EXP | 2.2 | 2.5E-10 | 1.8 | 1.5E-05 | |
| ***TMSB4X*** | Xp22.2 | thymosin beta 4, X-linked | GEN/EXP | 1.9 | 3.4E-04 | 2.0 | 1.5E-04 | Targets of *MYC* and *MAX* identified by ChIP on chip in a Burkitt's lymphoma cell line; overlap set.[9]<br>Genes down-regulated by *MYC*, according to the *MYC* Target Gene Database.[11] |
| ***COL9A2*** | 1p34.2 | collagen type IX alpha 2 chain | GEN/EXP | 2.4 | 3.4E-09 | 1.5 | 3.2E-02 | Genes directly down-regulated in P493-6 cells (B lymphocyte, Burkitt's lymphoma) by *MYC*.[8] |
| ***AFF2*** | Xq28 | AF4/FMR2 family member | GEN/EXP | 2.1 | 5.1E-06 | 1.8 | 1.9E-03 | |
| ***SGPP1*** | 14q23.2 | sphingosine-1-phosphate phosphatase 1 | GEN/EXP | 1.8 | 6.1E-08 | 2.0 | 5.3E-10 | |
| ***ADAM28*** | 8p21.2 | ADAM metallopeptidase domain 28 | GEN/EXP | 2.1 | 1.9E-06 | 1.7 | 2.9E-03 | |
| ***RGS13*** | 1q31.2 | regulator of G protein signaling 13 | GEN/EXP | 1.9 | 5.0E-03 | 1.9 | 3.9E-03 | |
| ***KIAA0408*** | 6q22.33 | KIAA0408 | GEN/EXP | 2.1 | 5.8E-14 | 1.7 | 8.7E-08 | |
| ***ZSCAN18*** | 19q13.43 | zinc finger and SCAN domain containing 18 | GEN/EXP | 1.9 | 1.7E-05 | 1.9 | 1.7E-05 | |
| ***CTHRC1*** | 8q22.3 | collagen triple helix repeat containing 1 | GEN/EXP | 1.9 | 2.3E-04 | 1.9 | 6.0E-04 | |
| ***MIR155HG*** | 21q21.3 | MIR155 host gene | GEN/EXP | 2.1 | 1.5E-04 | 1.6 | 3.9E-02 | Genes up-regulated by *MYC* and whose promoters are bound by *MYC*, according to *MYC* Target Gene Database.[11]<br>Genes whose promoters are bound by *MYC*, according to *MYC* Target Gene Database.[11] |
| ***COL24A1*** | 1p22.3 | collagen type XXIV alpha 1 chain | GEN/EXP | 1.7 | 8.7E-07 | 2.0 | 2.0E-10 | |
| ***FGF2*** | 4q28.1 | fibroblast growth factor 2 | GEN/EXP | 2.2 | 1.0E-08 | 1.5 | 1.6E-02 | |
| ***NRIP1*** | 21q11.2-q21.1 | nuclear receptor interacting protein 1 | GEN/EXP | 1.9 | 5.6E-10 | 1.8 | 5.8E-08 | |
| ***NR3C2*** | 4q31.23 | nuclear receptor subfamily 3 group C member 2 | GEN/EXP | 2.0 | 5.0E-14 | 1.7 | 6.2E-08 | |
| ***SV2C*** | 5q13.3 | synaptic vesicle glycoprotein 2C | GEN/EXP | 1.7 | 2.5E-04 | 2.0 | 4.8E-06 | |
| ***GBA3*** | 4p15.2 | glucosylceramidase beta 3 | GEN/EXP | 1.9 | 6.4E-04 | 1.8 | 1.5E-03 | |
| ***OSBPL1A*** | 18q11.2 | oxysterol binding protein like 1A | GEN/EXP | 2.1 | 3.0E-09 | 1.6 | 5.0E-04 | |
| ***VPREB3*** | 22q11.23 | V-set pre-B cell surrogate light chain 3 | GEN | 2.1 | 7.2E-05 | 1.5 | 5.2E-02 | |
| ***SLC40A1*** | 2q32.2 | solute carrier family 40 member | GEN/EXP | 2.0 | 4.6E-09 | 1.6 | 2.4E-04 | |

| | | | | | | | | | |
|---|---|---|---|---|---|---|---|---|---|
| ***PAX5*** | 9p13.2 | paired box 5 | GEN/EXP | 1.9 | 4.6E-04 | 1.7 | 1.0E-02 | | Genes directly down-regulated in P493-6 cells (B lymphocyte, Burkitt's lymphoma) by *MYC*.[8] |
| ***LINC00494*** | 20q13.13 | long intergenic non-protein coding RNA 494 | GEN/EXP | 1.7 | 1.1E-06 | 1.9 | 2.0E-08 | | |
| ***ATP10B*** | 5q34 | ATPase phospholipid transporting 10B (putative) | EXP | 1.6 | 5.1E-02 | 2.0 | 1.7E-03 | | |
| ***GNG2*** | 14q22.1 | G protein subunit gamma 2 | GEN/EXP | 1.6 | 6.9E-04 | 2.0 | 1.5E-07 | | Genes directly down-regulated in P493-6 cells (B lymphocyte, Burkitt's lymphoma) by *MYC*.[8] |
| ***RND3*** | 2q23.3 | Rho family GTPase 3 | GEN | 2.2 | 2.8E-04 | 1.4 | 1.8E-01 | YES[1] | |
| ***TNFSF8*** | 9q32-q33.1 | TNF superfamily member 8 | GEN/EXP | 1.9 | 1.9E-04 | 1.7 | 5.4E-03 | | |
| ***FRMPD3*** | Xq22.3 | FERM and PDZ domain containing 3 | GEN/EXP | 1.6 | 1.1E-02 | 1.9 | 4.9E-04 | | |
| ***LRP11*** | 6q25.1 | LDL receptor related protein 11 | GEN/EXP | 1.9 | 1.1E-08 | 1.6 | 8.5E-05 | | |
| ***ZCCHC2*** | 18q21.33 | zinc finger CCHC-type containing 2 | GEN/EXP | 1.9 | 5.6E-07 | 1.7 | 1.3E-04 | | |
| ***PTPRJ*** | 11p11.2 | protein tyrosine phosphatase, receptor type J | GEN/EXP | 1.8 | 2.1E-06 | 1.7 | 1.1E-04 | | |
| ***NEK6*** | 9q33.3 | NIMA related kinase 6 | GEN/EXP | 2.1 | 3.0E-08 | 1.5 | 1.3E-02 | YES[2] | |
| ***ALOX5AP*** | 13q12.3 | arachidonate 5-lipoxygenase activating protein | GEN/EXP | 1.7 | 6.6E-03 | 1.9 | 9.4E-04 | | Genes positively correlated with amplifications of *MYC* in small cell lung cancer cell lines.[17] |
| ***DMKN*** | 19q13.12 | dermokine | GEN/EXP | 1.5 | 6.1E-03 | 2.0 | 2.3E-07 | | |
| ***TIAM1*** | 21q22.11 | T-cell lymphoma invasion and metastasis 1 | GEN/EXP | 1.8 | 7.9E-06 | 1.7 | 2.0E-04 | | Wnt target genes up-regulated after Cre-lox knockout of *APC* in the small intestine that require functional *MYC*.[12] Genes that interact with *MYC* by Genomatix MatBase database of transcription factors.[18] |
| ***SH3TC1*** | 4p16.1 | SH3 domain and tetratricopeptide repeats 1 | GEN/EXP | 1.9 | 9.0E-08 | 1.6 | 3.2E-04 | | |
| ***ZYX*** | 7q34 | zyxin | GEN/EXP | 1.9 | 1.1E-07 | 1.6 | 1.1E-04 | | |
| ***PARM1*** | 4q13.3 | prostate androgen-regulated mucin-like protein 1 | GEN/EXP | 1.8 | 2.3E-10 | 1.6 | 7.7E-07 | | |
| ***WNT5B*** | 12p13.33 | Wnt family member 5B | GEN/EXP | 1.7 | 2.1E-06 | 1.8 | 5.5E-08 | | Genes down-regulated in primary epithelial breast cancer cell culture over-expressing *MYC* gene.[4] |
| ***B3GALNT1*** | 3q26.1 | beta-1,3-N-acetylgalactosaminyltransferase 1 (globoside blood group) | GEN/EXP | 1.8 | 9.4E-06 | 1.7 | 2.9E-04 | | |
| ***PTPRC*** | 1q31.3-q32. | protein tyrosine phosphatase, receptor type C | GEN/EXP | 1.9 | 1.2E-05 | 1.5 | 1.0E-02 | | Genes down-regulated in B cell lymphoma tumors expressing an activated form of *MYC*.[15] Genes down-regulated in K562 cells (lymphoblast, chronic myelogenous leukemia) by *MYC* in the presence of *CKN1B*.[7] Genes directly down-regulated in P493-6 cells (B lymphocyte, Burkitt's lymphoma) by *MYC*.[8] |
| ***LPGAT1*** | 1q32.3 | lysophosphatidylglycerol acyltransferase 1 | GEN/EXP | 1.8 | 2.0E-08 | 1.6 | 1.6E-05 | | |
| ***MIAT*** | 22q12.1 | myocardial infarction associated transcript (non-protein coding) | GEN/EXP | 1.6 | 2.3E-02 | 1.8 | 1.3E-03 | | |
| ***RIMS3*** | 1p34.2 | regulating synaptic membrane exocytosis 3 | GEN/EXP | 1.9 | 1.9E-10 | 1.4 | 2.2E-03 | | |
| ***FRMD6*** | 14q22.1 | FERM domain containing 6 | GEN/EXP | 1.9 | 1.1E-08 | 1.4 | 7.5E-03 | | |
| ***RASGRP3*** | 2p22.3 | RAS guanyl releasing protein 3 | GEN/EXP | 1.9 | 8.0E-11 | 1.4 | 2.7E-03 | | |
| ***CRIM1*** | 2p22.2 | cysteine rich transmembrane BMP regulator 1 | GEN/EXP | 1.4 | 2.9E-02 | 1.9 | 1.1E-05 | | |
| ***SOCS1*** | 16p13.13 | suppressor of cytokine signaling 1 | GEN/EXP | 1.9 | 4.5E-08 | 1.4 | 4.8E-03 | | Genes directly down-regulated in P493-6 cells (B lymphocyte, Burkitt's lymphoma) by *MYC*.[8] |
| ***NLGN4X*** | Xp22.32-p22.31 | neuroligin 4, X-linked | EXP | 1.5 | 7.3E-02 | 1.8 | 6.7E-03 | | |
| ***SLC44A2*** | 19p13.2 | solute carrier family 44 member 2 | GEN/EXP | 1.5 | 1.6E-03 | 1.8 | 3.9E-07 | | |
| ***BMP2K*** | 4q21.21 | BMP2 inducible kinase | GEN/EXP | 1.9 | 2.8E-11 | 1.4 | 3.2E-03 | YES[1,2] | |
| ***AHR*** | 7p21.1 | aryl hydrocarbon receptor | GEN/EXP | 1.8 | 4.9E-07 | 1.5 | 4.3E-03 | | Genes that regulate *MYC* by Genomatix MatBase database of transcription factors.[18] |
| ***DSG2*** | 18q12.1 | desmoglein 2 | EXP | 1.2 | 4.1E-01 | 2.0 | 1.8E-03 | | Genes directly up-regulated in P493-6 cells (B lymphocyte, Burkitt's lymphoma) by *MYC*.[8] |
| ***MYOF*** | 10q23.33 | myoferlin | GEN/EXP | 1.4 | 4.7E-02 | 1.9 | 1.0E-04 | | |

| | | | | | | | | | |
|---|---|---|---|---|---|---|---|---|---|
| ***GRASP*** | 12q13.13 | general receptor for phosphoinositides 1 associated scaffold protein | GEN/EXP | 1.8 | 7.3E-05 | 1.5 | 2.4E-02 | | |
| ***SYNPO*** | 5q33.1 | synaptopodin | GEN/EXP | 1.4 | 4.9E-03 | 1.8 | 1.6E-06 | | |
| ***WDFY3*** | 4q21.23 | WD repeat and FYVE domain containing 3 | GEN/EXP | 1.9 | 2.4E-06 | 1.4 | 2.3E-02 | | |
| ***TNFSF12*** | 17p13.1 | TNF superfamily member 12 | GEN/EXP | 1.8 | 1.1E-05 | 1.4 | 3.6E-02 | | |
| ***RRAS2*** | 11p15.2 | RAS related 2 | GEN | 2.0 | 1.7E-05 | 1.2 | 4.3E-01 | | |
| ***DOK4*** | 16q21 | docking protein 4 | GEN/EXP | 1.8 | 1.1E-11 | 1.4 | 2.0E-03 | | |
| ***BIRC3*** | 11q22.2 | baculoviral IAP repeat containing 3 | GEN | 2.1 | 5.1E-06 | 1.1 | 7.7E-01 | | Genes whose promoters are bound by *MYC*, according to *MYC* Target Gene Database.[11] |
| ***KCNN4*** | 19q13.31 | potassium calcium-activated channel subfamily N member 4 | GEN | 1.9 | 4.1E-04 | 1.3 | 3.2E-01 | YES[1,2] | Genes up-regulated hT-RPE cells (immortalized retinal pigment epithelium) by *MYC*.[19] |

***GEN/EXP**: gene was significantly de-regulated in cases with abnormal *MYC* genomic profile as well as in cases with *MYC* expression log2>=13.0 with fold-change >=1.8 at least in one of these two tested parameters. **GEN**: gene was significantly de-regulated in cases with abnormal *MYC* genomic profile with fold-change >=1.8 and not significant in cases with *MYC* expression log2>=13.0. **EXP**: gene was significantly de-regulated in cases with *MYC* expression log2>=13.0 with fold-change >1.8 and not significant in cases with abnormal *MYC* genomic profile

References:

1. Wingender E. The TRANSFAC project as an example of framework technology that supports the analysis of genomic regulation. Brief Bioinform. 2008;9(4):326-332.
2. Xie X, Lu J, Kulbokas EJ, et al. Systematic discovery of regulatory motifs in human promoters and 3' UTRs by comparison of several mammals. Nature. 2005;434(7031):338-345.
3. Liberzon A, Birger C, Thorvaldsdottir H, Ghandi M, Mesirov JP, Tamayo P. The Molecular Signatures Database (MSigDB) hallmark gene set collection. Cell Syst. 2015;1(6):417-425.
4. Bild AH, Yao G, Chang JT, et al. Oncogenic pathway signatures in human cancers as a guide to targeted therapies. Nature. 2006;439(7074):353-357.
5. Schlosser I, Holzel M, Hoffmann R, et al. Dissection of transcriptional programmes in response to serum and c-Myc in a human B-cell line. Oncogene. 2005;24(3):520-524.
6. Schuhmacher M, Kohlhuber F, Holzel M, et al. The transcriptional program of a human B cell line in response to Myc. Nucleic Acids Res. 2001;29(2):397-406.
7. Acosta JC, Ferrandiz N, Bretones G, et al. Myc inhibits p27-induced erythroid differentiation of leukemia cells by repressing erythroid master genes without reversing p27-mediated cell cycle arrest. Mol Cell Biol. 2008;28(24):7286-7295.
8. Zeller KI, Zhao X, Lee CW, et al. Global mapping of c-Myc binding sites and target gene networks in human B cells. Proc Natl Acad Sci U S A. 2006;103(47):17834-17839.
9. Ben-Porath I, Thomson MW, Carey VJ, et al. An embryonic stem cell-like gene expression signature in poorly differentiated aggressive human tumors. Nat Genet. 2008;40(5):499-507.
10. Fernandez PC, Frank SR, Wang L, et al. Genomic targets of the human c-Myc protein. Genes Dev. 2003;17(9):1115-1129.
11. Zeller KI, Jegga AG, Aronow BJ, O'Donnell KA, Dang CV. An integrated database of genes responsive to the Myc oncogenic transcription factor: identification of direct genomic targets. Genome Biol. 2003;4(10):R69.
12. Sansom OJ, Meniel VS, Muncan V, et al. Myc deletion rescues Apc deficiency in the small intestine. Nature. 2007;446(7136):676-679.
13. Lee JS, Chu IS, Mikaelyan A, et al. Application of comparative functional genomics to identify best-fit mouse models to study human cancer. Nat Genet. 2004;36(12):1306-1311.
14. O'Donnell KA, Yu D, Zeller KI, et al. Activation of transferrin receptor 1 by c-Myc enhances cellular proliferation and tumorigenesis. Mol Cell Biol. 2006;26(6):2373-2386.
15. Yu D, Cozma D, Park A, Thomas-Tikhonenko A. Functional validation of genes implicated in lymphomagenesis: an in vivo selection assay using a Myc-induced B-cell tumor. Ann N Y Acad Sci. 2005;1059(145-159.
16. Ceballos E, Munoz-Alonso MJ, Berwanger B, et al. Inhibitory effect of c-Myc on p53-induced apoptosis in leukemia cells. Microarray analysis reveals defective induction of p53 target genes and upregulation of chaperone genes. Oncogene. 2005;24(28):4559-4571.
17. Kim YH, Girard L, Giacomini CP, et al. Combined microarray analysis of small cell lung cancer reveals altered apoptotic balance and distinct expression signatures of MYC family gene amplification. Oncogene. 2006;25(1):130-138.
18. Genomatix Matrix Library v11.0 (www.genomatix.de)
19. Alfano D, Votta G, Schulze A, et al. Modulation of cellular migration and survival by c-Myc through the downregulation of urokinase (uPA) and uPA receptor. Mol Cell Biol. 2010;30(7):1838-1851.
